# Reduced functional fungal communities in two species of sloths (*Bradypus variegatus* and *Choloepus hoffmanni*) suggest a link to slow digestion

**DOI:** 10.1101/2024.07.19.604311

**Authors:** Priscila Chaverri, Efraín Escudero-Leyva, Darling Mora-Rojas, Andrea Calvo-Obando, Mariana González, Esteban Escalante-Campos, Esteve Mesén-Porras, Daniela Wicki-Emmenegger, Diego Rojas-Gätjens, Judith Avey-Arroyo, Mariana Campos-Hernández, Erick Castellón, Andrés Moreira-Soto, Jan Felix Drexler, Max Chavarría

## Abstract

Sloths, with their ruminant-like digestive systems, possess the slowest digestion among mammals due to their low metabolic rate, minimal food intake, and extremely low-energy diet. However, no comprehensive studies have characterized the sloth’s gut microbiota, including fungi, and their role in digestion. This study hypothesized that effective plant fiber-degrading fungi (e.g., Neocallimastigomycota) would be scarce in the sloth’s gut. The aim was to describe the gut microbiota of three-toed (*Bradypus variegatus*) and two-toed (*Choloepus hoffmanni*) sloths to understand their link to slow digestion. Microbial composition and functionality were analyzed using shotgun metagenomics, metatranscriptomics, fungal metabarcoding (ITS 1 and 2 nrDNA), and cellulose degradation analysis. Microbial communities were dominated by bacteria (92–97%), followed by viruses (1–7%). Fungi accounted for only 0.06–0.5% of metagenomic reads and 0.1% of transcripts. Functional analysis revealed minimal CAZy abundance (1.7–1.9% in metagenomes, 0.2% in metatranscriptomes), with no fungal CAZys or glycoside hydrolases detected. Neocallimastigomycota had negligible abundance in metagenomic data and was absent in metatranscriptomic or ITS metabarcoding data. *Bradypus variegatus* showed overall lower CAZy abundance and fungal presence compared to *Choloepus hoffmanni*. Lastly, cellulose degradation analyses revealed that only ∼5–35% of the intake was digested. This study highlights the unique microbial ecosystem in sloths’ guts, showing minimal presence of plant fiber-degrading anaerobic fungi and limited microbial CAZys, aligning with their slow digestion and low metabolic rate, thus enhancing our understanding of their digestive efficiency and metabolic adaptations.

## MAIN

Sloths (families Megatherioidea and Mylodontoidea, order Pilosa) are considered the herbivorous mammals with the slowest digestion^1,2^. The rate of passage has been estimated to vary between 11 to 30 days, with an average of 16 days^1^. Low metabolic rate, little food intake, and an extremely low-energy diet have been linked to a slow rate of digestion^1,3–5^. Sloths have a poikilothermic strategy to regulate their body temperature, which can range between ∼20 and 40 °C depending on ambient temperatures^6,7^. To lower their body temperature, sloths have to also lower their resting metabolic rate^6,7^. Other studies hypothesize that their slow digestion is associated with the tough cellulose/lignin and plant toxins that may require longer digestion periods and detoxification^8–11^. However, to date, there are no comprehensive studies that have characterized the sloth’s gut microbiome and its potential role in digestion. Out of the limited published research, one study showed that the rate of fermentation associated with gut bacteria in the three-toed sloth *Bradypus tridactylus* was significantly slower than that reported in other small foregut fermenters^12^. Another study revealed that the three-toed sloth (*Bradypus variegatus*) had reduced gut bacterial communities compared to the two-toed sloth (*Choloepus hoffmanni*)^13^. None of those studies considered fungi or viruses.

Given the markedly slower rate of digesta passage and defecation frequency observed in wild and captive sloths compared to ruminants and other herbivores (approximately 5–20 times slower)^14–16^, the present study hypothesized that the abundance of effective plant fiber-degrading taxa, such as anaerobic fungi (AF) (e.g., Neocallimastigomycota), would be notably scarce within the sloth’s gut microbiota. As a herbivore, it would be expected that the sloth’s gut microbiome resembled that of other herbivores, especially mammals with complex digestive system anatomies^17,18^. Sloths are foregut fermenters and polygastric, with seven gastric compartments resembling ruminant stomachs^19^. Studies have shown that ruminant or ruminant-like microbiomes are dominated by bacteria, followed by eukaryotes (fungi and protozoans) and archaea^17,18,20–23^.

Even though some estimate that fungi constitute only a small portion of the total microbial biomass in most herbivores (∼1–20%)^20,22,24^, these microorganisms are essential in the breakdown, digestion, and fermentation of complex plant polysaccharides^20,21,25–28^. Some have even suggested that the contribution of AF to the degradation of plant material could be more significant than the contribution of cellulolytic bacteria^21,29–31^. Although rumen fungal zoospore numbers are low in comparison to counts of bacteria and archaea, AF have been shown to comprise up to 20% of the rumen microbial biomass^32^ and account for 10–16% of rRNA transcript abundance^22^. Qi et al.^33^ also found that fungal reads dominated the ruminant’s gut metatranscriptome (∼16%), whereas 7% belonged to Alveolata, 5% to Metazoa, and 3.5% to Bacteria; >60% of the total reads were unclassified. Fungal cellulases generally account for >10% of the total glycoside hydrolases (GH) cellulases in the rumen, the majority produced by the AF phylum Neocallimastigomycota^20,21,26,28^.

The main objective of this study was to characterize the gut microbiota, with a specific focus on fungi, in both three-toed (*B. variegatus*) and two-toed (*C. hoffmanni*) sloths, aiming to shed light on the relationship with their slow digestion and metabolism. Fecal samples from individuals kept in a sloth sanctuary were collected. Microbiota profiling (metagenomes and metatranscriptomes, and fungal metabarcoding) was done to first determine the diversity of microbial communities, with emphasis on fungi, and then to infer if the functionality of the microbiota (e.g., enzymes involved in breaking down plant polysaccharides) is another factor contributing to the slow digestion and metabolism in sloths. Lastly, cellulose content before (plant diet) and after (fecal samples) passage through the sloth’s digestive tract was assessed to verify if the plant material was poorly digested. The results of this study will contribute to a better understanding of the role of fungi (and other microorganisms) in the unique digestive processes and metabolic adaptations in sloths. In addition, by providing foundational evidence on the role of microorganisms in the slow digestion and metabolism characteristic of sloths, the present study aims to deepen insights into these charismatic mammals’ biology, ecology, evolution, and conservation efforts.

## RESULTS

The relative abundances of the different microbial groups were first estimated in fecal samples from the two sloth species using shotgun metagenomics and metatranscriptomics. Using Kaiju or Kraken taxonomy classifiers, it was revealed that bacteria dominated the classified gut microbiota (∼92–97%), followed by viruses (∼1–7%); fungi accounted for only 0.06–0.5% of metagenomic reads and 0.1% of transcripts (Figure 1; Supplementary Figures S1 and S2 and Tables S1–S7). The fungal reads consisted of an absence or negligible abundance of the AF group Neocallimastigomycota in metagenomic and metatranscriptomic data (Figure 2; Supplementary Figure S3 and Tables S1–S7). For example, in *B. variegatus*, the abundance of AF was ∼5% of the fungal metagenome reads and 0.003% of the total microbial biomass. The abundance of AF in *C. hoffmanni* was ∼16% of the fungi, and 0.006% of the total microbial metagenome reads. The fungal phyla that dominated fecal samples were Ascomycota (∼30–40% of the total fungal reads), Mucoromycota (∼16–30% of the total fungal reads), and Basidiomycota (∼16–22%) (Figure 2; Supplementary Figure S3 and Tables S1–S7). Fungal transcripts from *C. hoffmanni* were also dominated by Mucoromycota (∼30%; e.g., *Rhizopus*), followed by Ascomycota (∼38%; e.g., *Saccharomycodes*) (Supplementary Tables S6 and S7). With ITS nrDNA metabarcoding, it was further confirmed that Neocallimastigomycota or other fungi that may aid in the breakdown of the complex plant polysaccharides were either absent or present in negligible amounts, as no ITS sequences attributable to this group were detected (Supplementary Figures S4–S6 and Tables S8 and S9). The most abundant fungal taxa found, e.g., Saccharomycetes, followed by Sordariomycetes, Dothideomycetes, and Eurotiomycetes in the Ascomycota, belong to groups that are not known for their effective fibrolytic role in herbivore guts^31,34^.

**Figure 1.**
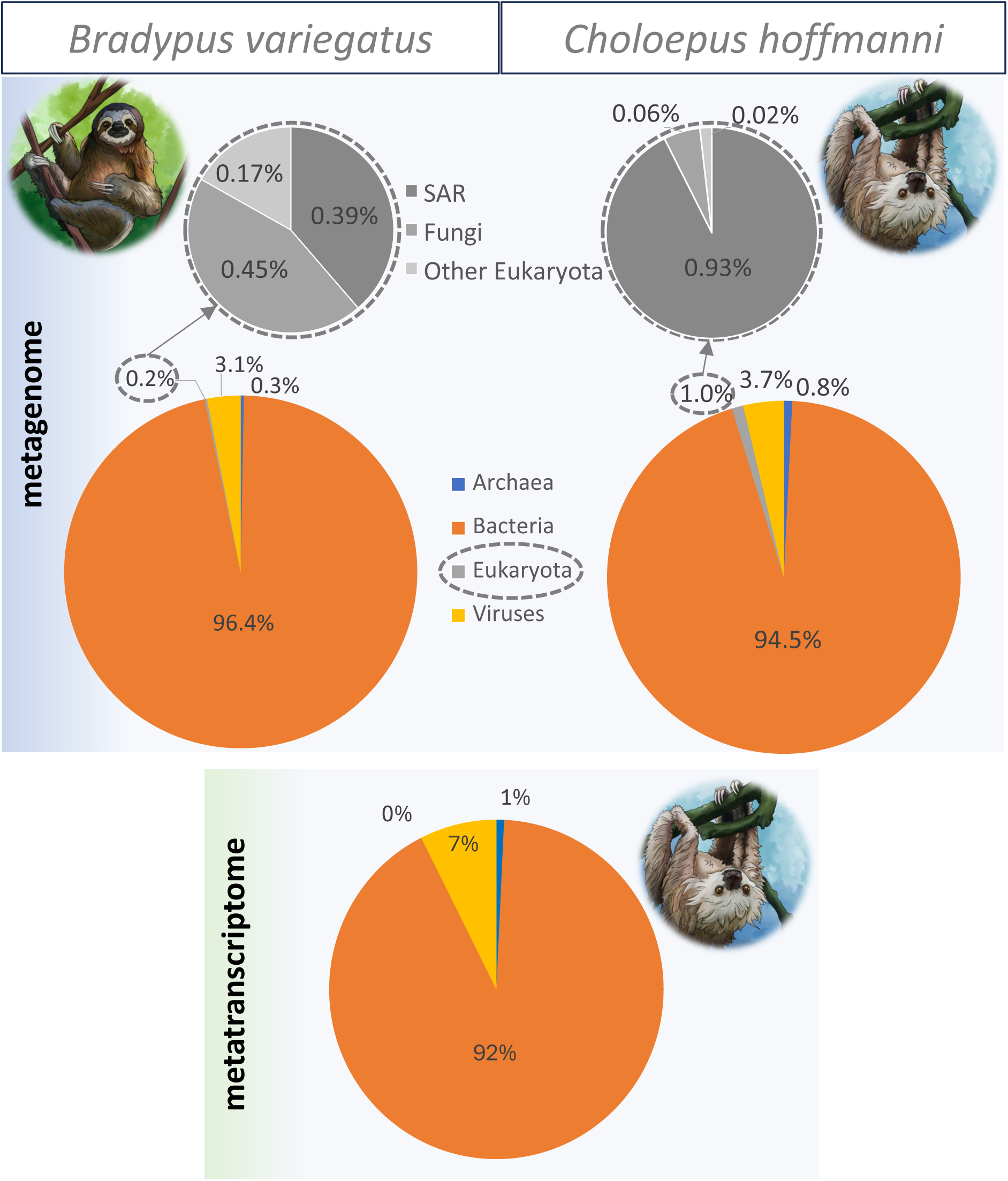
Relative abundance (%) of classified microbial groups from two species of sloths. SAR supergroup includes stramenopiles, alveolates, and Rhizaria. Relative abundances presented here are based on the results from the search with Kaiju taxonomy classifier. For *B. variegatus*, 55.4% were unclassified and 19.6% could not be assigned to a non-viral species. For *C. hoffmanni*, 37.6% were unclassified and 23% could not be assigned to a non-viral species.

**Figure 2.**
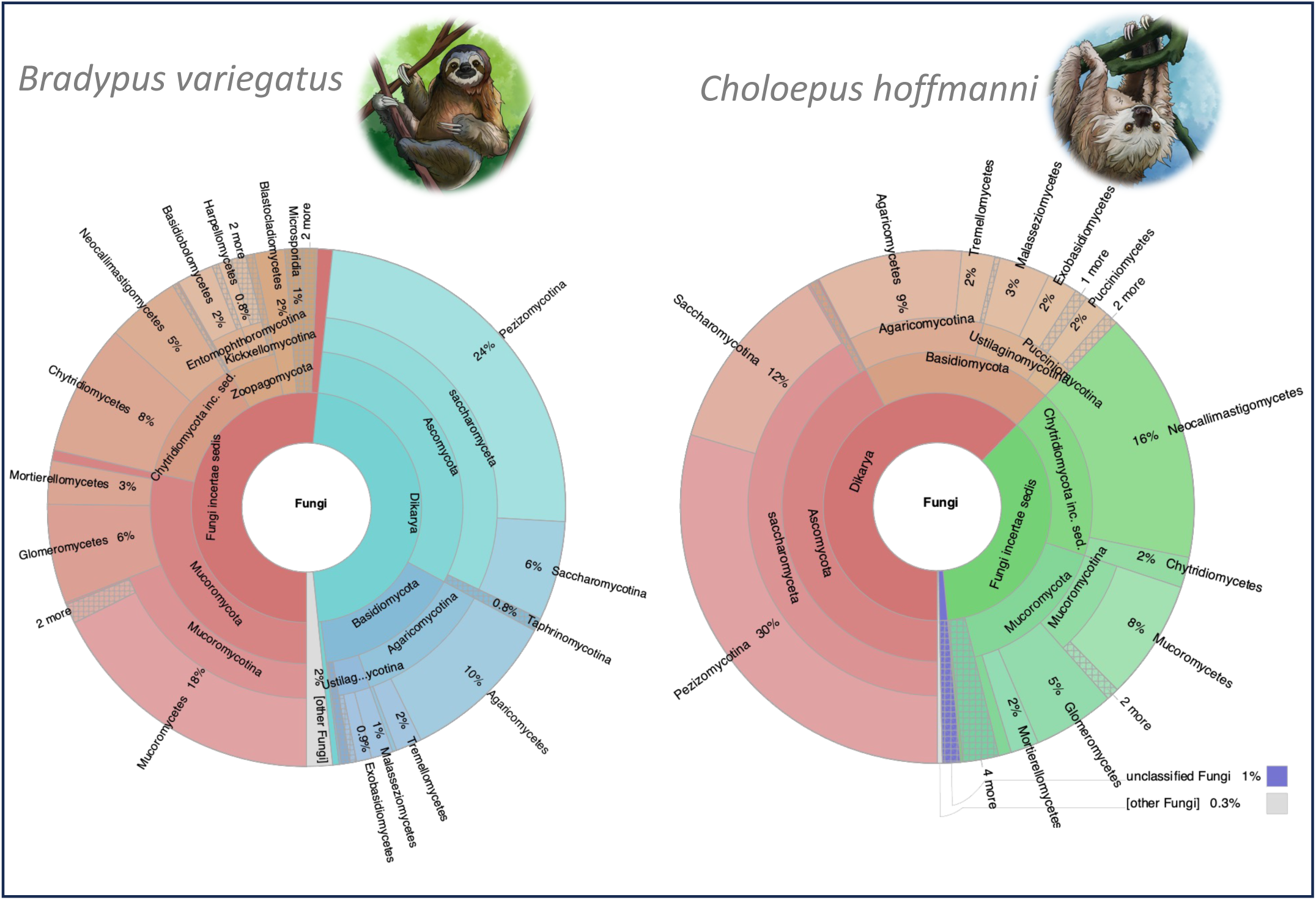
Relative abundances (%) of <u>fungal</u> taxa in two species of sloths. Relative abundances presented here are based on the results from the search with Kaiju taxonomy classifier and metagenomic data. Unclassified reads are not included. Interactive Krona figures in HTML format are included in https://github.com/pchaverri/sloths. These HTML files include all taxa (Prokaryota, Eukaryota and Viruses).

The bacterial taxa Bacillota (=Firmicutes; e.g., Bacilli and Clostridia), Bacteroidetes (e.g., Bacteroidia), and Proteobacteria were the most prevalent, constituting 33–48%, 9–20%, and 5– 24%, respectively (Figure 3; Supplementary Figure S7 and Tables S1 and S2). At lower taxonomic levels, Oscillospiraceae (∼12–21%; e.g., *Oscillibacter*, *Pseudoflavonifractor*, and *Ruminococcus*); Lachnospiraceae (∼12–15%; e.g., *Lachnoclostridium* and *Roseburia*); and Clostridiaceae (∼8%; e.g., *Clostridium*) dominated the bacterial communities (Supplementary Figure S7 and Tables S1 and S2). Actinobacteria was the fourth most abundant bacterial phylum in the present study (∼2– 5% of bacteria; Supplementary Figure S7 and Tables S1 and S2). This phylum was dominated by members of *Corynebacterium*, followed by *Bifidobacterium* and *Gulosibacter* (Supplementary Table S2).

**Figure 3.**
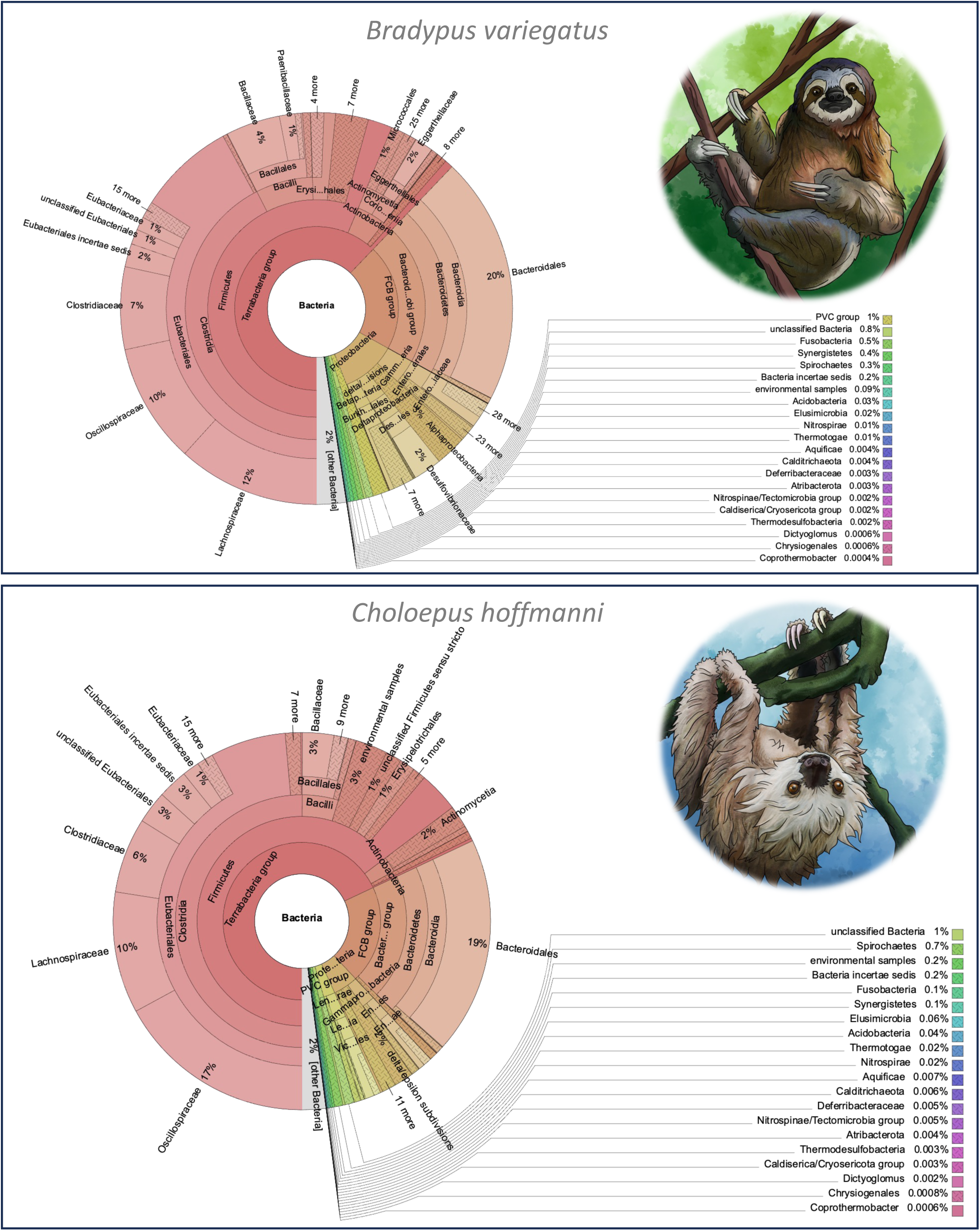
Relative abundances (%) of <u>bacterial</u> taxa in two species of sloths. Relative abundances presented here are based on the results from the search with Kaiju taxonomy classifier and metagenomic data. Unclassified reads are not included. Interactive Krona figures in HTML format are included in https://github.com/pchaverri/sloths. These HTML files include all taxa (Prokaryota, Eukaryota and Viruses).

On the other hand, expected taxa, such as *Fibrobacter* and *Prevotella*, were in very low abundance (0.01–0.3% and 0.1–1.4%, respectively); the former was absent from transcripts, and the abundance of *Prevotella* was ∼0.5% (Supplementary Tables S1, S2, S6, and S7). Archaea accounted for 0.3–0.8% of the classified microbial reads, and ∼46–78% of the total archaeal reads were classified as Methanobacteria. Transcripts from *C. hoffmanni* samples were dominated by Bacillota (∼50%; e.g., *Pseudoflavonifractor, Streptococcus, Clostridium, Blautia*, and *Mediterraneibacter*), followed by Proteobacteria (∼19%; e.g., *Escherichia*, *Klebsiella*, and *Vibrio*), Actinobacteria (∼7.5%), and Bacteroidetes (∼7.5%; e.g., *Bacteroides*) (Supplementary Tables S6 and S7).

The beta diversity analysis revealed that the overall microbial communities of both species of sloths are different, exhibiting heterogeneity among the species (p = 0.008598) and significance for the dispersion (p = 0.0002). Visualization through Non-metric Multidimensional Scaling (NMDS) supports these differences (stress value = 0.0150; Supplementary Fig. S8). Linear Discriminant Analysis (LDA) also revealed that the key microbial taxa that differentiate the microbial communities in the two species of sloths belong to bacterial groups. Several noteworthy examples are Coriobacteriia, Eggerthellales, Rhodobacterales, and Proteobacteria in *B. variegatus*; and *Bifidobacterium,* Clostridiales, *Flavonifractor*, *Oscillibacter*, *Pseudoflavonifractor,* and *Ruminococcus* in *C. hoffmanni* (Supplementary Fig. S9). *Bifidobacterium*, *Ruminococcus, Pseudoflavonifractor,* and *Oscillibacter* were 7×, 5×, 4×, 4×, respectively, more abundant in *C. hoffmanni* than in *B. variegatus* (Supplementary Table S2).

Clusters of orthologous groups (COGs) belonging to carbohydrate-active enzymes (CAZy) were characterized from the fecal metagenomes and metatranscriptome using eggNOG-mapper. CAZys are central in breaking down complex carbohydrates, such as cellulose, lignin, and xylan^35^, and fungi are central producers of these enzymes^27,36,37^. Functional analysis revealed minimal CAZy abundance (1.7–1.9% in metagenomes, 0.2% in metatranscriptome), with no fungal CAZys detected in the *C. hoffmanni* metatranscriptome. Shotgun metagenomic results showed that CAZy COGs were in low abundance (1.7–1.9% of the total number of predicted COGs) (Table 1 and Supplementary Tables S10–14). The great majority of the CAZys matched to bacteria (i.e., Clostridia and Bacteroidetes) (Figure 4 and Supplementary Tables S10–14). No archaeal, fungal, or other eukaryotic CAZys were detected (or were insignificant) in *B. variegatus*; and only 0.009% corresponded to non-bacterial CAZys in *C. hoffmanni* (Table 1). Eleven (65%) of the fungal CAZys from *C. hoffmanni* were from Ascomycota and six (35%) from Basidiomycota (Figure 4 and Supplementary Tables S10–S12). Only 55 CAZy OGs were detected in the metatranscriptomic data (Supplementary Tables S13 and S14). Glycoside hydrolases (GH), which are essential CAZys for the processing of plant polysaccharides^35,38^, were detected mostly from bacteria in both sloth species (339 in *B. variegatus* and 10,100 in *C. hoffmanni*); a few were identified from fungi (12) and other eukaryotes (17) only in *C. hoffmanni* (Table 1) in the metagenomic data. 27 GHs were detected in the metatranscriptome, all corresponding to bacteria (e.g., Bacilli, Bacteroidetes, and Gammaproteobacteria) (Supplementary Tables S13 and S14). No fungal GH transcripts were detected. Moreover, there was a negligible abundance (0–0.1% of the total microbiota) of auxiliary-active enzymes (AAs) (Table 1), which are also fundamental in delignification of plant biomass^39^.

**Figure 4.**
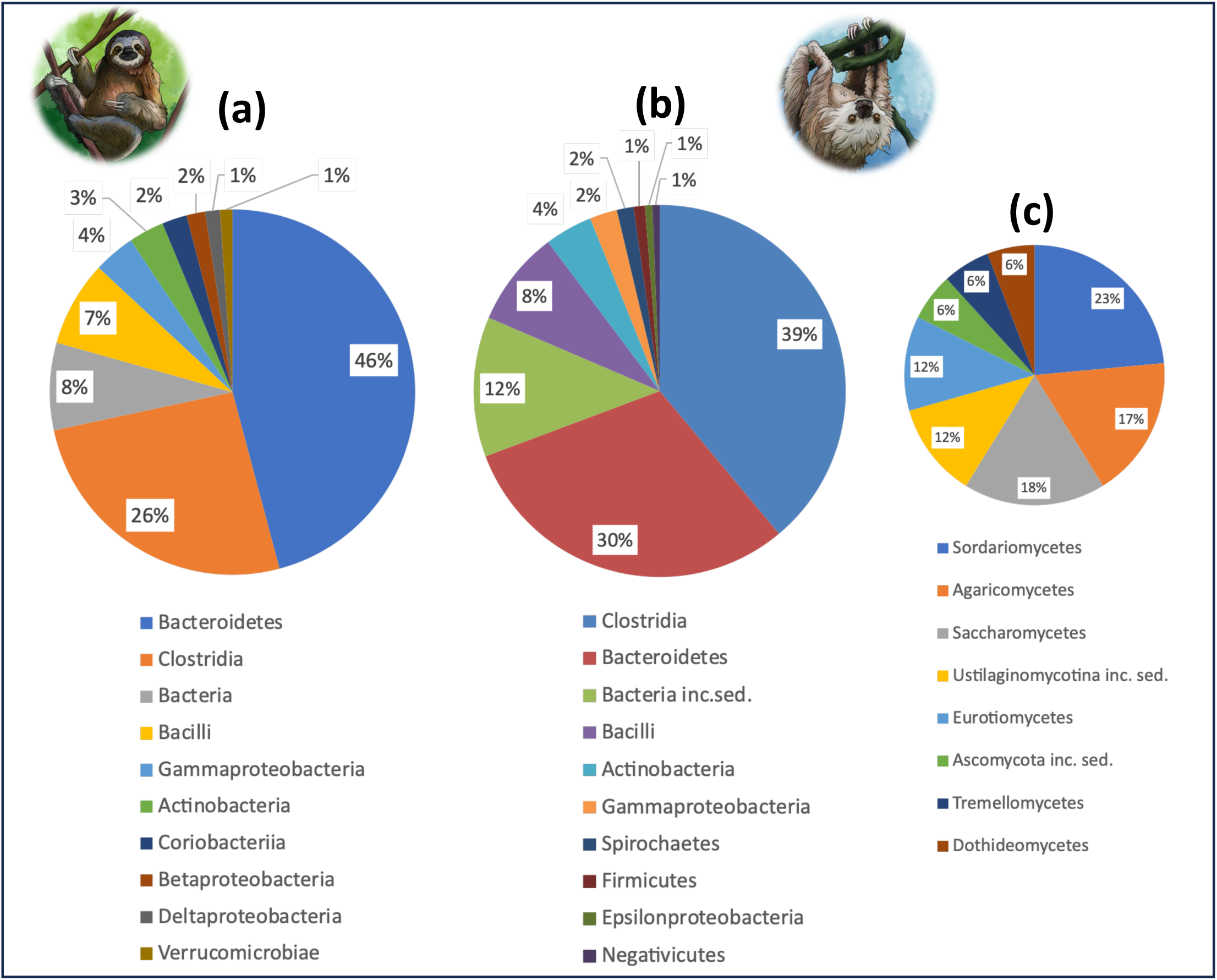
Relative abundance (%) of carbohydrate-active enzymes (CAZy) from shotgun metagenomic data for each sloth species and bacterial and fungal group. Only top-ten most abundant groups are shown. (a) *Bradypus variegatu*s bacterial CAZys. (b) *Choloepus hoffmanni* bacterial CAZys. (c) *C. hoffmanni* fungal CAZys. No fungal CAZys were detected in *B. variegatus*.

**Figure 5.**
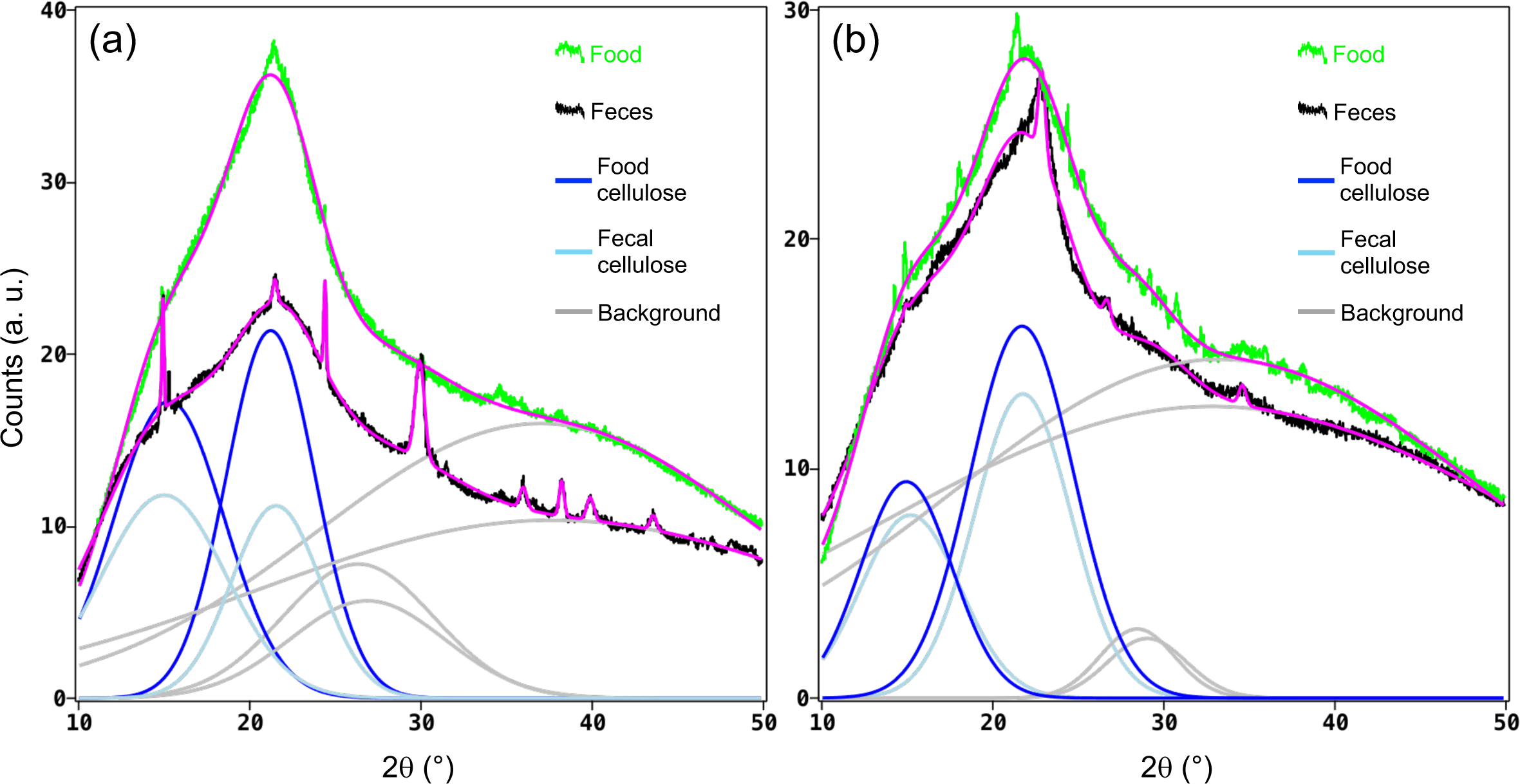
X-ray diffractograms of powders of food and the corresponding feces for sloth species *B. variegatus* (a) and *C. hoffmanni* (b), for one replicate. Green noisy curves correspond to the experimental signals of foods, black noisy curves correspond to experimental fecal signals. Blue and light blue curves are deconvoluted Gaussian functions assigned to cellulose in the food and feces, respectively. Gray curves represent background Gaussian functions. Magenta curves represent the fitted functions which are the sum of the background functions and the deconvoluted peaks assigned to cellulose. The sharp peaks present in the diffractogram of *Bradypus* feces are due to calcium oxalate crystals (RRUFF Database, https://rruff.info/Whewellite/R050526); several Gaussian peaks were added to conveniently fit these signals but were excluded from the figure for the sake of simplicity. X-ray diffractograms for all three replicates are in Supplementary Figure S10.

**Table 1.**
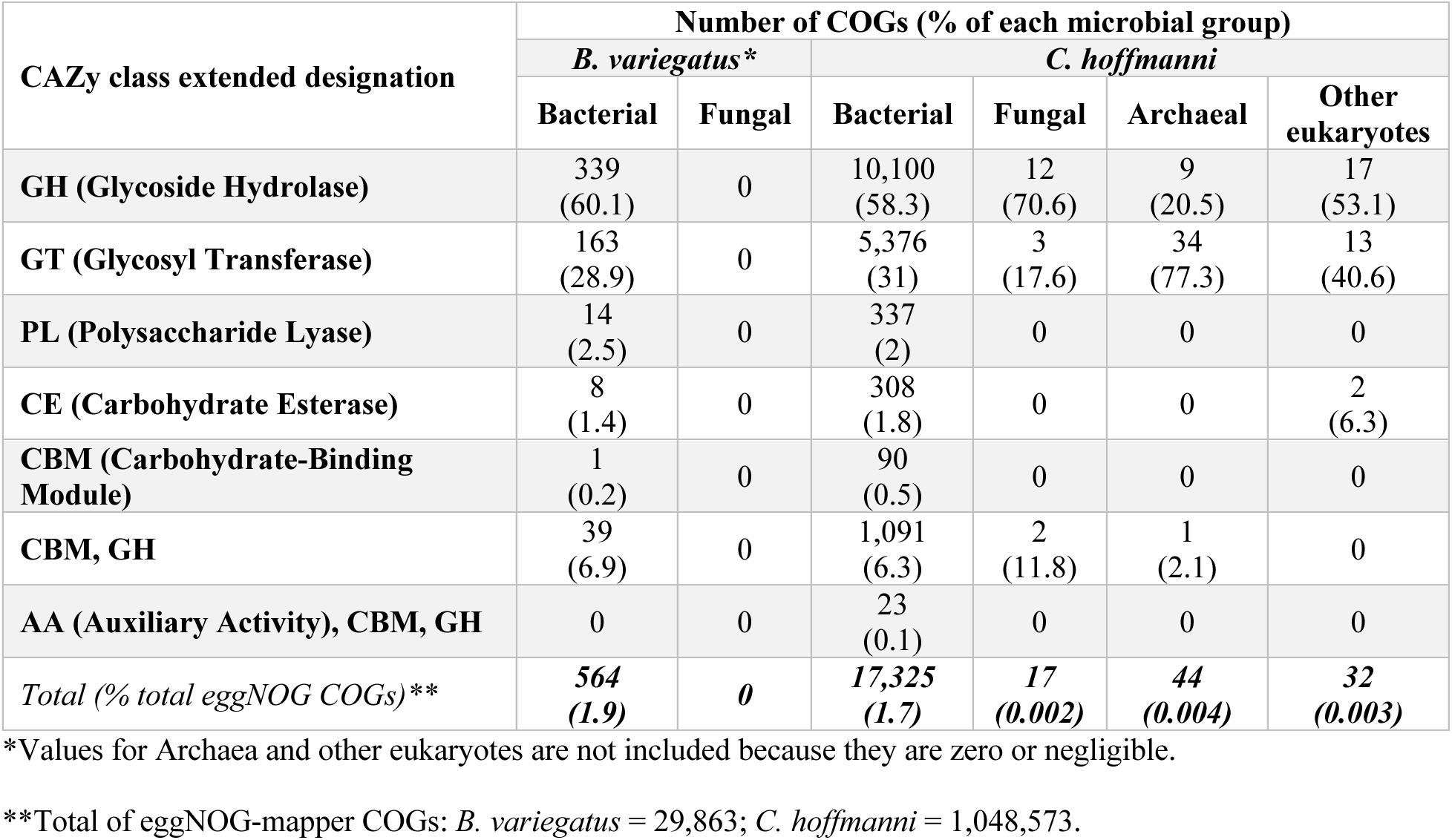
Designation of shotgun metagenomic eggNOG clusters of orthologous groups (COGs) into carbohydrate-active enzyme (CAZy) classes for each sloth species and microbial group. See supplementary Tables S10–12 for data used to construct this table.

| CAZy class extended designation | Number of COGs (% of each microbial group) |  |  |  |  |  |
| --- | --- | --- | --- | --- | --- | --- |
|  | *B. variegatus* * |  | *C. hoffmanni* |  |  |  |
|  | Bacterial | Fungal | Bacterial | Fungal | Archaeal | Other eukaryotes |
| **GH (Glycoside Hydrolase)** | 339<br>(60.1) | 0 | 10,100<br>(58.3) | 12<br>(70.6) | 9<br>(20.5) | 17<br>(53.1) |
| **GT (Glycosyl Transferase)** | 163<br>(28.9) | 0 | 5,376<br>(31) | 3<br>(17.6) | 34<br>(77.3) | 13<br>(40.6) |
| **PL (Polysaccharide Lyase)** | 14<br>(2.5) | 0 | 337<br>(2) | 0 | 0 | 0 |
| **CE (Carbohydrate Esterase)** | 8<br>(1.4) | 0 | 308<br>(1.8) | 0 | 0 | 2<br>(6.3) |
| **CBM (Carbohydrate-Binding Module)** | 1<br>(0.2) | 0 | 90<br>(0.5) | 0 | 0 | 0 |
| **CBM, GH** | 39<br>(6.9) | 0 | 1,091<br>(6.3) | 2<br>(11.8) | 1<br>(2.1) | 0 |
| **AA (Auxiliary Activity), CBM, GH** | 0 | 0 | 23<br>(0.1) | 0 | 0 | 0 |
| *Total (% total eggNOG COGs)\*\** | **564**<br>**(1.9)** | **0** | **17,325**<br>**(1.7)** | **17**<br>**(0.002)** | **44**<br>**(0.004)** | **32**<br>**(0.003)** |

\*Values for Archaea and other eukaryotes are not included because they are zero or negligible.
\*\*Total of eggNOG-mapper COGs: *B. variegatus* = 29,863; *C. hoffmanni* = 1,048,573.

From the results of eggNOG-mapper, the presence and abundance of specific key enzymes involved in lignocellulolytic (e.g., laccase, lignin peroxidase, manganese peroxidase, and versatile peroxidase; CAZy families AA1 and AA2); cellulolytic (e.g., endoglucanase, exoglucanase, β-D glucosidase, and cellobiose dehydrogenase; GH3, GH5, GH6, GH7, GH9, GH48, CBM1, CBM2, AA3, AA7, and AA8); hemicellulolytic (e.g., endoxylanase, arabinofuranosidase, esterase, and α-glucuronidase; CBM1, CBM2, CBM13, CBM22, CBM72, GH5, GH8, GH10, GH11, and GH67, among others); and xylanolytic (e.g., xylanases; GH10, GH11, and GH30, among others) activity^38^ were inspected. From the metagenomic data, lignocellulolytic enzymes were absent across all fecal microbiomes examined (Supplementary Tables S10–14). On the other hand, β-glucosidases (GH3; EC 3.2.1.21) were the most abundant (e.g., GH3 = ∼11% and 10% of the CAZys in *C. hoffmanni* and *B. variegatus*, respectively); only one CAZy from the GH3 family matched to fungi. GH3 family had only two transcripts. GH13 and GH77 families, which contain enzymes involved in degradation of starches and malto-oligosaccharides (e.g., α-amylases)^40^, were also abundant CAZys in metagenome data (∼14% and 17% of the total GHs in *B. variegatus* and *C. hoffmanni*, respectively); ∼60% from the total GH transcripts of *C. hoffmanni* were GH13+GH77. Three GH13 COGs were from fungi in *C. hoffmanni* and none from *B. variegatus*. Exo- and endoglucanases (e.g., GH5 and GH9) were also present in CAZy metagenome data (∼9% and 4% of the total CAZys in *C. hoffmanni* and *B. variegatus*, respectively), but most were from bacteria; only two were from fungi in *C. hoffmanni* (Supplementary Tables S10–14). Other key enzymes involved in plant fibrolysis were in very low abundance or absent. For example, β-xylanases (e.g., GH30; EC 3.2.1.8; 1 COG in *B. variegatus* and 49 in *C. hoffmanni*, all from bacteria).

Finally, a comparative analysis of the cellulose content in both the sloths’ dietary intake and fecal samples was performed using X-ray diffraction (XRD), used as an indicator of the general degradation of plant biomass. The comparison of corresponding deconvoluted peaks assigned to cellulose in feces and food revealed a degradation ratio of 30±4% for *B. variegatus* and 11±7% for *C. hoffmanni* (Figure 4 and Supplementary Figure S10).

## DISCUSSION

The present study reveals that fungi have little to negligible abundance and function in the guts of two species of sloths. The limited fungal communities were dominated by Ascomycota, followed by Basidiomycota, and Mucoromycota, similar to those reported in other herbivore guts except ruminants or ruminant-like mammals . However, the most noteworthy result, which supports the original hypothesis of this study, is that the AF group Neocallimastigomycota, abundant in herbivore, ruminant, and ruminant-like guts (8–20% of the total microbial biomass)^20,21,26,32,42–44^, was insignificant or absent in metagenomic, metatranscriptomic, and metabarcoding data. In addition, shotgun metagenomic results showed that microbial (prokaryotes and eukaryotes) CAZy COGs, including glycoside hydrolases (GHs), were in very low abundance, 5–15× less than reported for other ruminant or ruminant-like gut microbiomes^39,45–48^. No fungal GH transcripts were detected and the great majority of the CAZys matched bacteria (i.e., Clostridia and Bacteroidetes). Moreover, the overall reduced abundance of auxiliary-active enzymes (AAs) adds to the apparent limited capability within the sloth microbiota for the delignification of plant biomass^39^.

Bacterial communities dominated the fecal samples (92–97%). The three most prevalent bacterial phyla found in the fecal samples (i.e., Bacillota, Bacteroidetes, and Proteobacteria) were the same as those reported by Dill-McFarland et al.^13^ in wild two- and three-toed sloths from Costa Rica, and by Delsuc et al.^49^ in fecal samples of two-toed sloths in captivity, although in different proportions. While Dill-McFarland et al.^13^ established that the dominant phyla in wild sloths were Proteobacteria and Bacillota, Delsuc et al.^49^ reported Bacteroidetes and Bacillota as the dominant phyla, similar to other mammalian herbivores. Dill-McFarland et al.^13^ speculated that the divergences in bacterial communities could be due to differences in diet between wild and captive animals. Other phyla found in samples from the present study are the same found in wild sloths^13^ (i.e., Verrucomicrobia, Fusobacteria, Lentisphaerae, Planctomycetes, Spirochaetes, and Tenericutes, among others). On the other hand, it was expected to find in more abundance other core bacterial constituents of herbivore and ruminant guts (e.g., *Fibrobacter*, Micrococcales, Micromonosporales, *Prevotella*, and Pseudonocardiales, among others)^33,50,51^. However, these were in very low abundance or absent from metagenomic or metatranscriptomic reads, aligning with findings from Dill-McFarland et al.^13^

Several abundant bacterial taxa identified, such as *Clostridium*, *Corynebacterium*, *Oscillibacter*, *Pseudoflavonifractor*, and *Ruminococcus*, are well-known residents of herbivore guts. These bacteria play crucial roles in fermenting and breaking down plant polysaccharides and detoxifying plant alkaloids and other secondary metabolites^17,42,50,52–55^. For instance, Lachnospiraceae (Eubacteriales), one of the most abundant families found in the present study, is a group of obligately anaerobic bacteria that are among the most abundant taxa in the rumen^50^ and in wild sloths^13^. Within the Eubacteriales, *Ruminococcus* spp. have been reported as a vital part of the gut microbiota in animals, humans, and the rumen^56–58^. In addition, consistent with reports from wild sloths^13^, Actinobacteria was the fourth most abundant bacterial phylum in the present study. Noteworthy examples are *Flavonifractor* and *Pseudoflavonifractor*. These genera are known for their ability to degrade flavonoids^52,53^, abundant in many plants, which may thus play a role in the detoxification of these compounds. Therefore, considering the low to negligible abundance of cellulose-degrading fungi (e.g., Neocallimastigomycota), some of the abovementioned bacterial taxa may be the main actors in plant biomass degradation in both captive and wild sloths.

The functional annotation results suggest that in both sloth species, a few enzymes, such as β-glucosidases (e.g., GH3), α-amylases (e.g., GH13 and GH77), and, to a lesser extent, exo- and endoglucanases (e.g., GH5 and GH9) from bacteria, are primarily responsible for breaking down cellulose and starch in both species of sloths (no lignocellulolytic enzymes were found). However, other important fibrolytic enzymes seem scarce. Endoglucanases, exoglucanases, and β-glucosidases need to act synergistically and simultaneously to break down cellulose through hydrolysis^31,39,59^. Fungi appear to potentially contribute only about 4–11% of these enzymes, as indicated by metagenome data; negligible contribution is shown in the metatranscriptome data. Therefore, the potential contribution of fungal enzymes in *B. variegatus* and *C. hoffmanni* guts to plant fibrolysis seems extremely limited.

X-ray diffraction (XRD) results align with the notion of reduced cellulose digestion in these animals. Specifically, *B. variegatus*, which is exclusively fed *Cecropia* spp. leaves, provide a simpler and more conclusive experimental system compared to *C. hoffmanni* sloths, whose mixed diet complicates analysis and interpretation. By comparing the cellulose content in *Cecropia* leaves and feces, it was determined that *B. variegatus* exhibits a cellulose degradation rate of 30±4%, significantly lower than the 50–80% reported for other herbivores, particularly ruminants^60–63^. In contrast, the results for *C. hoffmanni* are less definitive due to their mixed diet, which includes other macromolecules and nutrients (e.g., carrots contain approximately 1% protein and 4–5% sugars^64^), other than crude fibers, that are preferentially utilized as carbon sources by the animal and its microbiota. Consequently, the low cellulose degradation rate observed in *C. hoffmanni* (11±7%) may be attributed to their captive diet rather than an inherent lower cellulose degradation capacity.

The results of the present study may support a relationship but cannot unequivocally conclude that the reduced microbiota functionality is the main cause of a low metabolic rate and, consequently, slow digestion. This study shows significant differences in community composition and functionality between the sloths’ microbiotas; *B. variegatus* had less microbial CAZys (i.e., GH enzymes) and less potential abundance of Neocallimastigomycota fungi than *C. hoffmanni*. Pauli et al.^4^ found that *B. variegatus* had ∼30% lower field metabolic rate (FMR) and moved less than *C. hoffmanni*, *B. variegatus* being the mammal with the lowest FMR recorded for any mammal. Most studies suggest that gut microbiota influences various aspects of host metabolism, including energy extraction from food and regulation of host metabolism^65–67^. Gut microbes play a crucial role in fermenting indigestible carbohydrates and producing short-chain fatty acids, which can influence host metabolism, energy balance, and overall homeostasis^68^. Additionally, the gut microbiota can affect the absorption of nutrients and regulate the production of hormones involved in metabolism, such as insulin and glucagon^69,70^.

The low cellulose and digesta degradability observed in sloths, the negligible content of Neocallimastigomycota fungi, and the reduced functionality of the gut microbiota suggest a limited ability of sloths to extract energy from their diet, which may then result in a slow metabolism. While most studies suggest that gut microbiota influence host metabolism, others propose that the metabolic rate of the host and other host environmental factors can shape the composition and function of the gut microbiota^71–74^. Therefore, the relationship between gut microbiota and metabolic rate in mammals is complex and likely bidirectional^65–67,75^. For example, the host’s genetics, metabolism, diet, or other factors (e.g., immune function and gut physiology) can select for specific microbial species composition or influence the establishment and maintenance of the gut microbiota^74,76^. Undoubtedly, differences in dietary habits and metabolic efficiency among mammalian species can also shape the gut microbiota to optimize energy extraction from their specific diet^77,78^. *Bradypus variegatus* is an obligatory folivore (e.g., *Cecropia* spp.), while *C. hoffmanni* can live on a more varied diet of fruits, flowers, and leaves^2,4,13^. Dill-McFarland et al.^13^ had already found a relationship between the sloths’ diet and bacterial communities. They showed that *C. hoffmanni* had a richer gut bacterial community than that of *B. variegatus*, and that feeding behavior selected for specific microbial communities. Consistent with these findings, research has shown reduced fermentation activity in the digestive systems of *Bradypus tridactylus*^12,79^. This has been linked, at least in part, to the elevated lignin levels found in *Cecropia palmata*^79^. Consequently, this slower fermentation process affects the breakdown of leaf fibers and the detoxification of secondary compounds^10^. The findings of the present study, which include diminished fungal communities and functionality within the gut microbiotas, alongside elevated levels of undigested plant fibers, provide further support to conclusions from the abovementioned studies.

In conclusion, the present study revealed, for the first time, the extremely limited fungal presence (e.g., Neocallimastigomycota) and functionality in the sloths’ gut microbiotas, conflicting with what previous studies found in ruminants or ruminant-like mammals. For example, essential enzymes for plant polysaccharide breakdown primarily come from bacteria, while fungal contribution is minimal. While the present study does not conclusively link reduced microbiota functionality to low metabolic rates, differences between sloth species suggest potential impacts. *Bradypus variegatus* shows lower overall CAZy abundance and fungal (i.e., Neocallimastigomycota) presence compared to *Choloepus hoffmanni*, possibly influencing their metabolic rates. The results of this study provide fundamental knowledge to better understand the role of fungi (and other microorganisms) in the unique digestive processes and metabolic adaptations in sloths. Overall, the relationship between gut microbiota and metabolic rate in mammals involves a complex interplay of multi-directional host-microbe interactions that certainly deserve further scrutiny.

## METHODS

### Study site and sample collection

To generate less disturbance and discomfort to the animals, a non-invasive sampling method was used to analyze their gut microbiota (i.e., fecal samples). Feces were collected from three-toed (*Bradypus variegatus*) and two-toed (*Choloepus hoffmanni*) sloths kept at The Sloth Sanctuary (Limón, Costa Rica, 9°47’58.5"N 82°54’53.1"W). The sloths kept in the Sanctuary have been rescued from the wild. Their diets are strictly controlled, aiding in data standardization by reducing variability stemming from unknown dietary factors^80^. In addition, because their defecation events are sporadic, by having a controlled diet, data reproducibility is increased from fecal samples collected during different times and individuals^80^. In summary, implementing a controlled diet aimed to minimize the influence of external variables, including environmental exposures, lifestyle factors, and dietary variations. The diet of *B. variegatus* is based exclusively on *Cecropia* spp. leaves and that of *C. hoffmanni* is more varied (see Extended Methods for more detail). Due to the small number of animals and low defecation frequency, fewer samples were obtained from *B. variegatus* than for *C. hoffmanni* (see next section: “DNA and RNA extractions, and sequencing”). Samples for DNA extraction were collected, immediately transported on ice to the laboratory, and frozen at -80°C until processing. Samples for RNA extraction were collected and immediately suspended in RNAlater® buffer (Sigma Aldrich, USA). The samples were transported on ice to the laboratory, and frozen at -80°C until processing (see Extended Methods).

### DNA and RNA extractions, and sequencing

For ITS metabarcoding analysis, fecal samples from 3 *B. variegatus* and 8 *C. hoffmanni* individuals were used. For shotgun metagenomics fecal samples from 5 *B. variegatus* and 5 *C. hoffmanni* individuals were used. DNA was extracted using DNeasy PowerSoil kit (QIAGEN, Hilden, Germany), with modifications (see Extended Methods). To obtain a comprehensive microbiota profiling (e.g., prokaryotes, eukaryotes, and viruses), genomic DNA samples were sent to Novogene Inc. (Sacramento, CA, USA) for shotgun metagenome sequencing (Illumina NovaSeq PE150). To better characterize the fungal communities in the fecal samples, targeted amplicon metagenomics (metabarcoding) was also outsourced to Novogene (Illumina NovaSeq PE250). The Internal Transcribed Spacers of the nuclear ribosomal DNA (ITS nrDNA) region is considered the official barcode for fungi and thus, has the most comprehensive and curated databases^81,82^. For metabarcoding, ITS1 and ITS2 were sequenced to recover as much fungal diversity as possible^83^.

For metatranscriptomic analysis, 23 fecal samples from *C. hoffmanni* individuals were used. Unfortunately, due to their scarce defecation frequency, not enough *B. variegatus* fecal material was collected to obtain RNA with the quantity and quality for metatranscriptome sequencing. RNA extraction and metatranscriptome sequencing were performed at the sequencing facility of the German Centre for Infection Research (DZIF), Associated Partner Site Charité (Berlin, Germany). Fecal samples from *C. hoffmanii* kept in RNAlater buffer were transported to DZIF where total nucleic acids (RNA) were extracted using the MagNA Pure 96 DNA and Viral NA Large Volume Kit (Roche Diagnostics, Indianapolis, IN, USA) in a Magna Pure 96 Instrument (Roche Diagnostics), following the manufacturer’s protocols. The RNA extractions were pooled to have enough concentration and quality for sequencing. RNA libraries were prepared for the sample according to KAPA HyperPrep manufacturer protocol (Roche Diagnostics) and sequenced in an Illumina NextSeq 550 system (150 cycles paired-end).

### Bioinformatic analyses

#### Metagenomics, metatranscriptomics, and functional annotation

All bioinformatic analyses were run on the Kabré supercomputer (CNCA-CONARE, Costa Rica). Once the raw .fastq files were obtained from Novogene and DZIF, low-quality reads were filtered and trimmed with Seqtk v.1.4 (https://github.com/lh3/seqtk/) and BBDuk v.38.84 (https://sourceforge.net/projects/bbmap/), respectively (see Extended Methods for more detail). To remove host reads from the trimmed files, two available sloth genomes (*Bradypus variegatus* GCA_004027775.1 and *Choloepus hoffmanni* GCA_000164785.2) were used to create separate databases with those genomes using Bowtie2 v.2.2.9^84^. Following, the metagenome reads were mapped to the host databases using Bowtie2 and removed from further analysis. The assembly of the metagenomes or metatranscriptomes was done with SPAdes v.3.14.1 (“meta” or “rna”)^85–87^. EggNOG-mapper v.2.1.6^88^ was used for functional annotation of the assembled metagenome and metatranscriptome reads. EggNOG-mapper predicts orthologous groups and phylogenies from the eggNOG database (http://eggnog5.embl.de) to then transfer functional information from those predicted orthologous groups^88^. EggNOG-mapper was run using MMseqs as the search step algorithm^89^. The raw .fastq files have been deposited in GenBank under BioProject PRJNA991536.

Two tools to taxonomically classify the reads, i.e., Kaiju v.1.9.2^90^ and Kraken2^91^, were utilized and compared. For this, only the assembled contigs/scaffolds and transcripts that were longer than 500 bp were used. Kaiju has been reported as equally sensitive to Kraken2, but Kaiju is the only classifier that includes fungal and other eukaryotes databases from the NCBI RefSeq database and a protein-based classifier^90,92,93^. For Kaiju, the analysis was first run using “nr_euk” database v.2022-03-10, which includes prokaryotes, viruses, and eukaryotes. Because with nr_euk few reads matched fungi, the analysis was also run using only the fungi database v.2023-05-06. The analyses were run for each sample separately. For Kraken2, the standard database (v.2023-4-25), plus fungi and protozoans, was used. The results shown in the present study are based on Kaiju nr_euk database because when the search was run with the Kaiju fungi and Kraken2 databases, many fungal groups were missing (e.g., Neocallimastigomycota from Kaiju fungi, and most non-Dikarya from Kraken2) (Supplementary Table S1). Relative abundance (%) calculations and bar graphs were done in the package microeco v.1.1.0^94^ in RStudio v.2023.03.1+446. All the resulting and supplementary data are deposited in the public repository GitHub https://github.com/pchaverri/sloths.

#### Targeted amplicon metagenomics/metabarcoding of fungi

DADA2 v.1.26.0^95^ implemented in RStudio was utilized for processing raw sequencing data. This included quality inspection, filtering, trimming, merging of paired-end reads, inference of Amplicon Sequence Variants (ASVs)^96^, and subsequent removal of chimeras (adapters and primers were already removed by Novogene Inc.) (see Extended Methods for more detail). Taxonomic assignments were done by performing similarity searches against the curated and quality-checked UNITE database v.2021^82,97^ using the DECIPHER package v.2.0^98^. Relative abundance (%) calculations and bar/bubble graphs were also done in microeco and ggplots. The raw .fastq files have been deposited in GenBank under BioProject PRJNA991536. Additional results and supplementary data are deposited in GitHub public repository https://github.com/pchaverri/sloths.

#### Community analysis

The microbial communities of *B. variegatus* and *C. hoffmanni* were compared using shotgun metagenomic sequencing results. For beta diversity analysis, the Hellinger transformation was applied to the ASV counts table. The "adonis" and "permutest" functions from the Vegan package^99^ were used to evaluate differences in the microbial communities between sloth species and homogeneity across samples, with permutation parameters set to 5,000 for both tests. Visualization was performed using Non-metric Multidimensional Scaling (NMDS) in Phyloseq from Bioconductor^100^. For this, count data was transformed into a Bray-Curtis distance matrix, and the analysis was conducted with a set seed. Linear Discriminant Analysis (LDA) diagrams were created using the microeco package. Relative abundance, NMDS, LDA plots, and all analyses were performed using R v.4.3.3.

#### Comparative analysis of cellulose degradation using X-ray diffraction (XRD)

XRD was selected as the method for determining plant biomass degradation due to its ability to provide detailed structural information, quantify crystallinity, and monitor changes in a non-destructive and sensitive manner^101^. XRD of the samples was used to quantitatively assess crystalline and amorphous components present in materials^102,103^. The areas or intensities of deconvoluted signals in diffractograms assigned to a specific substance are considered proportional to the content of the substance. In the case of samples containing cellulose, the relative contents of crystalline and amorphous cellulose have been previously assessed with this deconvolution technique^104,105^. Identical procedures of sample preparation and XRD measurements were applied to all samples. Sloth food samples (*Cecropia* spp. leaves for *B. variegatus* and mixture for *C. hoffmanni*) and feces from both sloth species were prepared accordingly (see Extended Methods for more detail) to be analyzed with a Bruker D8 Advance ECO X-ray powder diffraction equipment with Bragg-Brentano geometry (specific conditions are detailed in Extended Methods). Each resulting diffractogram was deconvoluted by summing four Gaussian functions centered at 2θ values about 15°, 22°, 28° and 38°. Three samples of each sloth’s specific feed were analyzed at 2θ values between 10° and 50°, where signals associated with cellulose appear^106^. The first two Gaussian peaks were assigned to cellulose^106^, and the remaining were used to fit the background (signals assigned to other substances than cellulose, such as lignin, are considered as part of the background). The experimental diffractograms were fitted by non-linear regression using the software for symbolic and numeric computation Maple (Maplesoft, Waterloo, Canada). The areas of the peaks centered about 15° (𝐴15°) and 22° (𝐴22°) for the fecal samples were compared with the peak areas calculated for the corresponding food to estimate the cellulose degradation ratio for both species of sloths:

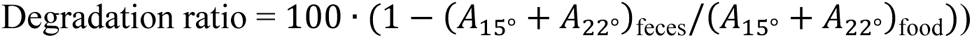

## Data Availability Statement

Metagenomic, metatranscriptomic, and metabarcoding data have been deposited in GenBank under BioProject PRJNA991536. Additional resulting and supplementary data are deposited in GitHub public repository https://github.com/pchaverri/sloths.

## Acknowledgments

The authors thank Noelia Rechnitzer, Angélica Chavarría and Juliana Chavarría for their assistance in one of the sampling campaigns.

## Author Contributions

PC and MC conceived and designed the experiments; EE-L, MG, DM-R, AC-O, EE-C, EM-P, DW-E, DR-G, and AM-S performed the experiments; PC, EE-L, EM-P analyzed the data; PC, JA-A, JFD, and MC acquired funds; PC, EE-L and MC wrote the paper. All authors reviewed and approved the final version of the manuscript.

---

**Figure S1.**
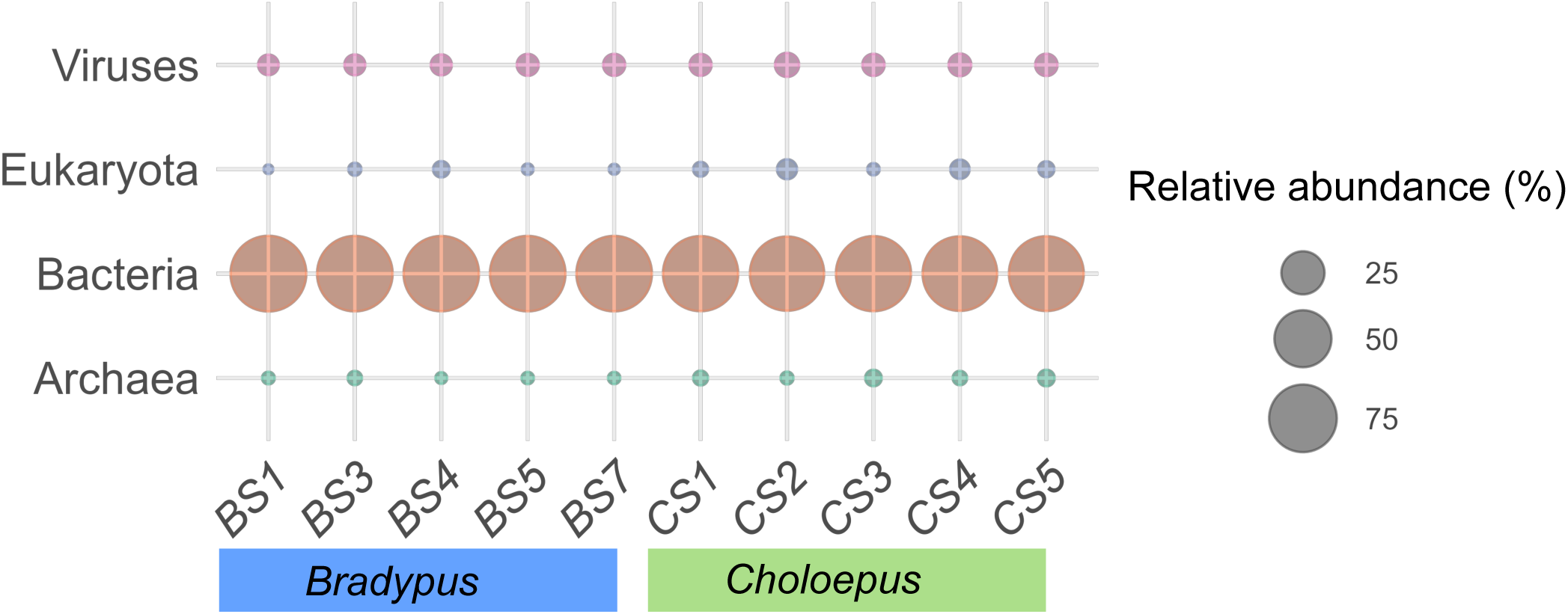
Relative abundance (%), per sample, of classified microbial groups from two species of sloths at the superkingdom/domain level. BS and CS represent samples from *B. variegatus* and *C. hoffmanni*, respectively. Unclassified reads are not included.

**Figure S2.**
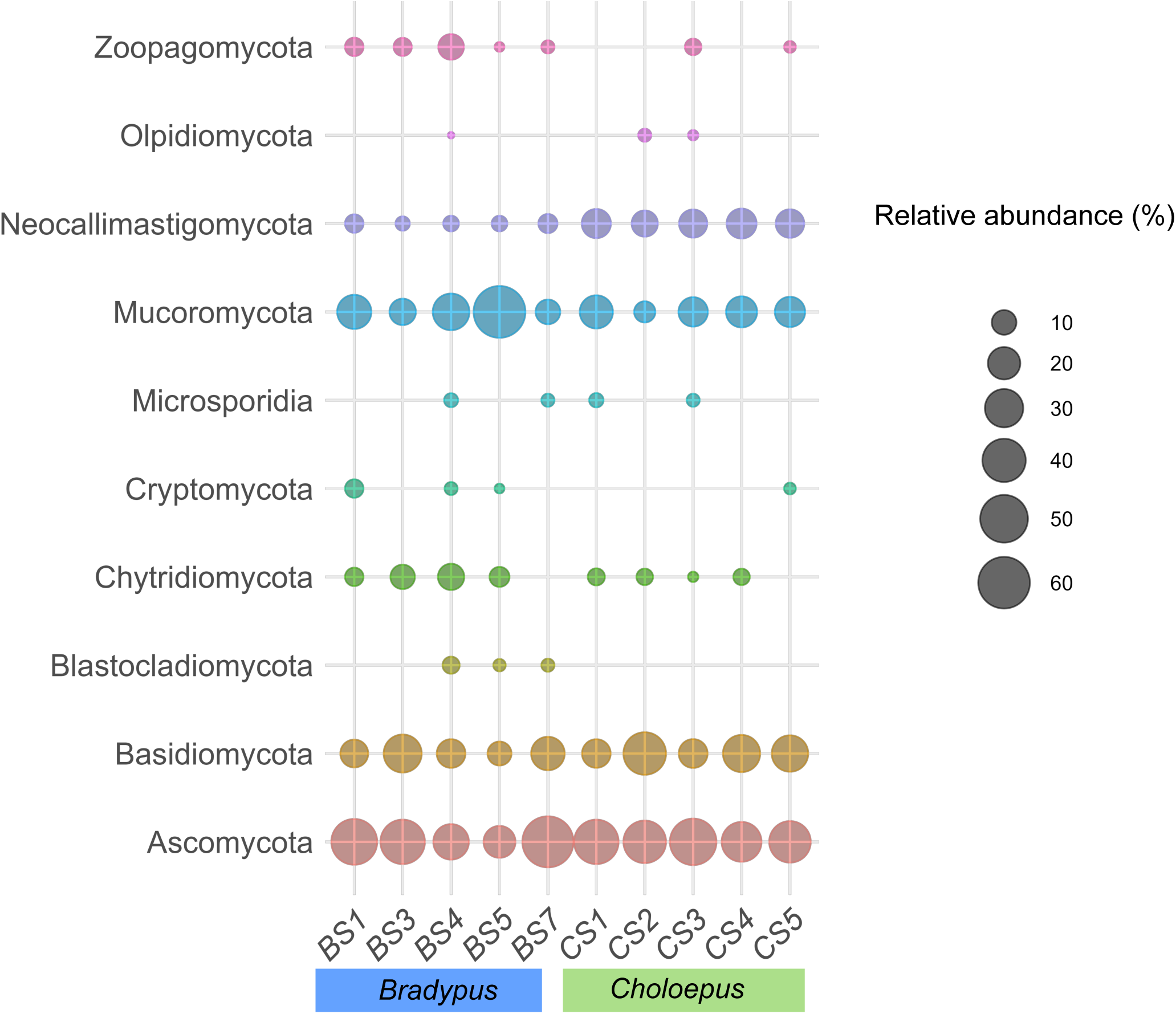
Relative abundance (%), per sample, of classified eukaryotes from two species of sloths at the kingdom level. BS and CS represent samples from *B. variegatus* and *C. hoffmanni*, respectively. Unclassified reads are not included. Results are based on metagenomic data.

**Figure S3.**
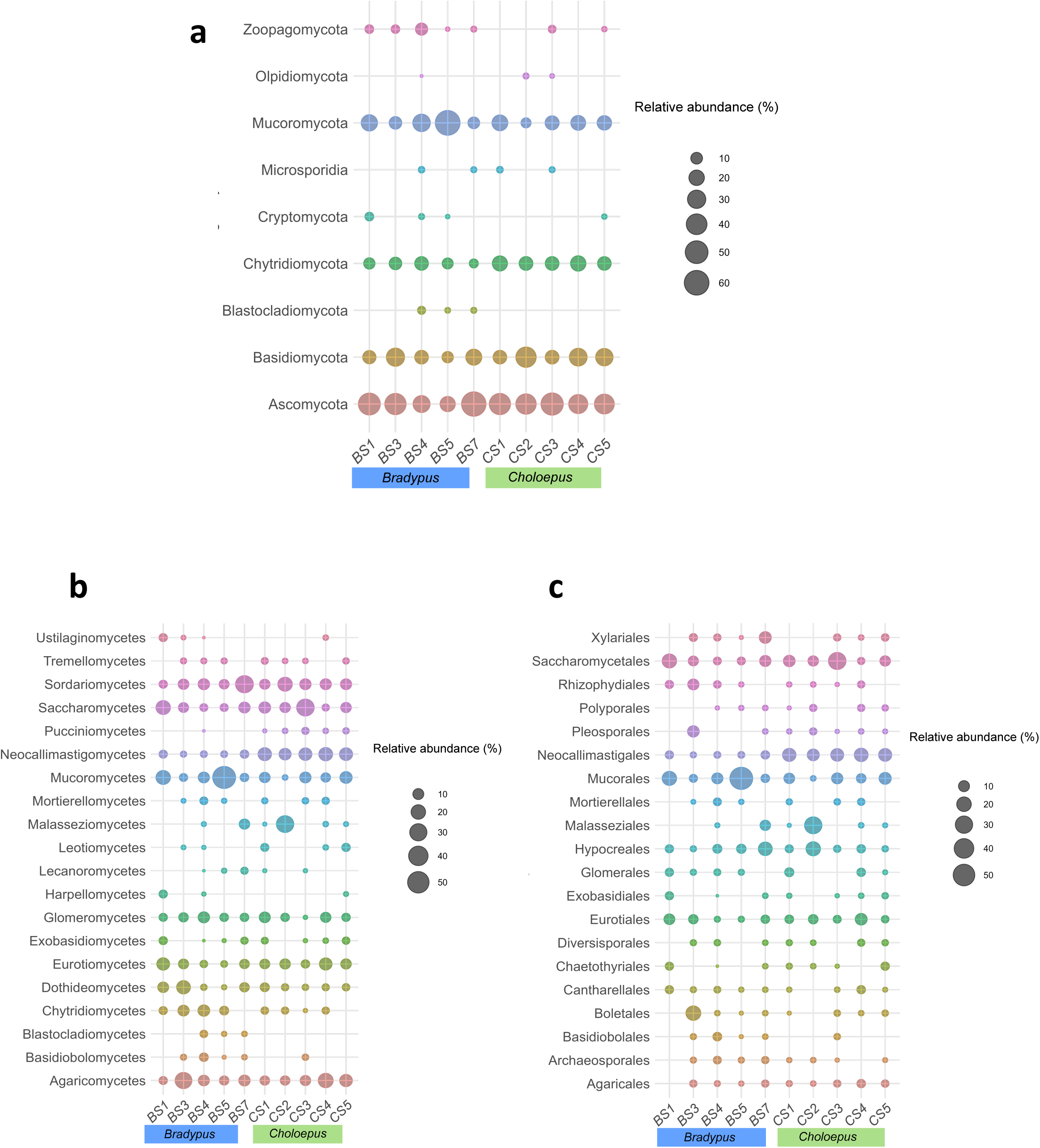
Relative abundances (%) of <u>fungal</u> taxa per sample in two species of sloths. Relative abundances presented here are based on the results from the search with Kaiju taxonomy classifier and metagenomic data. Fungal phyla (**a**), classes **(b),** and orders **(c)** per sample. Only the top 20 most abundant taxa are included. BS and CS represent samples from *B. variegatus* and *C. hoffmanni*, respectively. Unclassified reads are not included.

**Figure S4.**
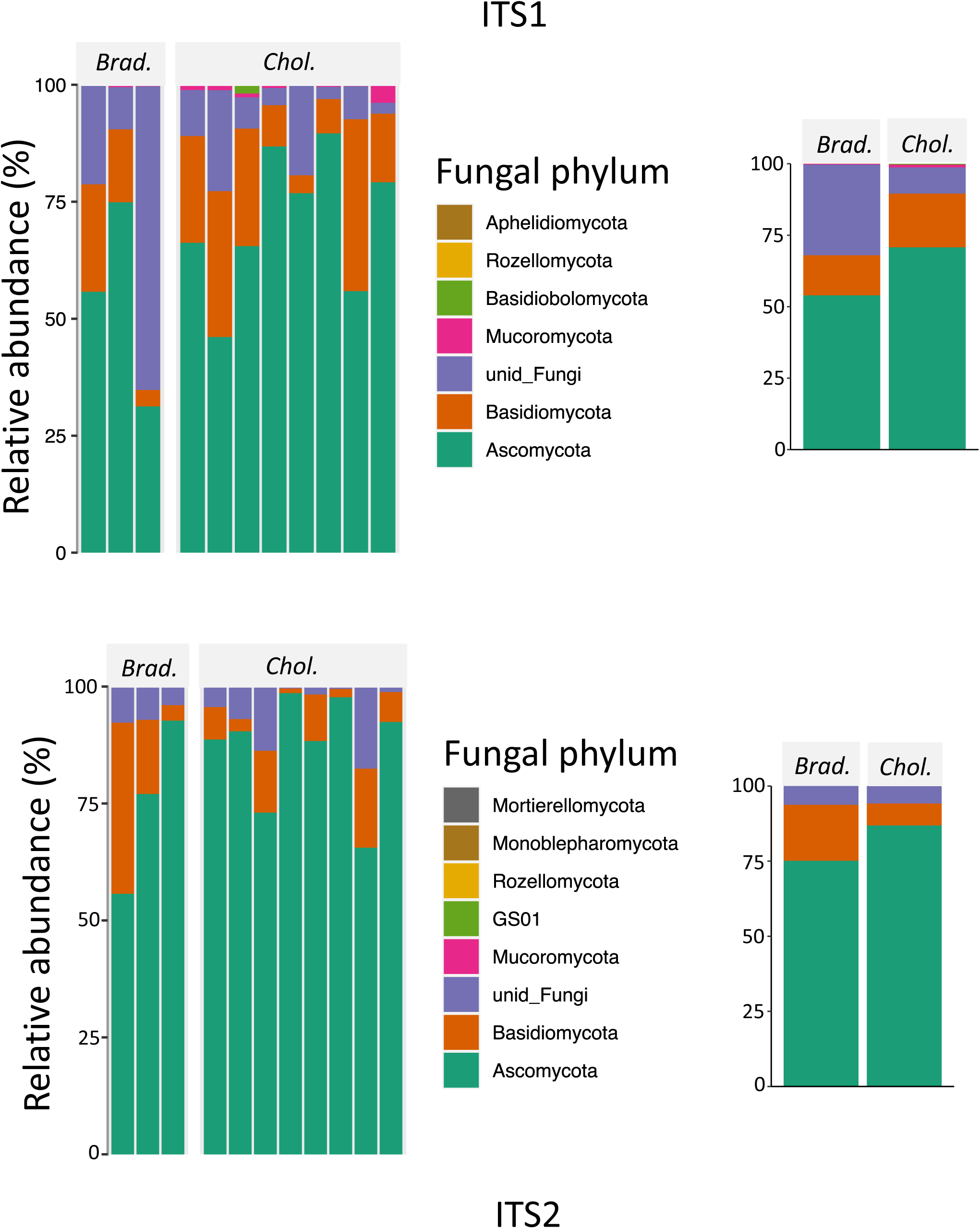
Relative abundance (%) of fungal phyla for ITS1 and ITS2 metabarcoding.

**Figure S5.**
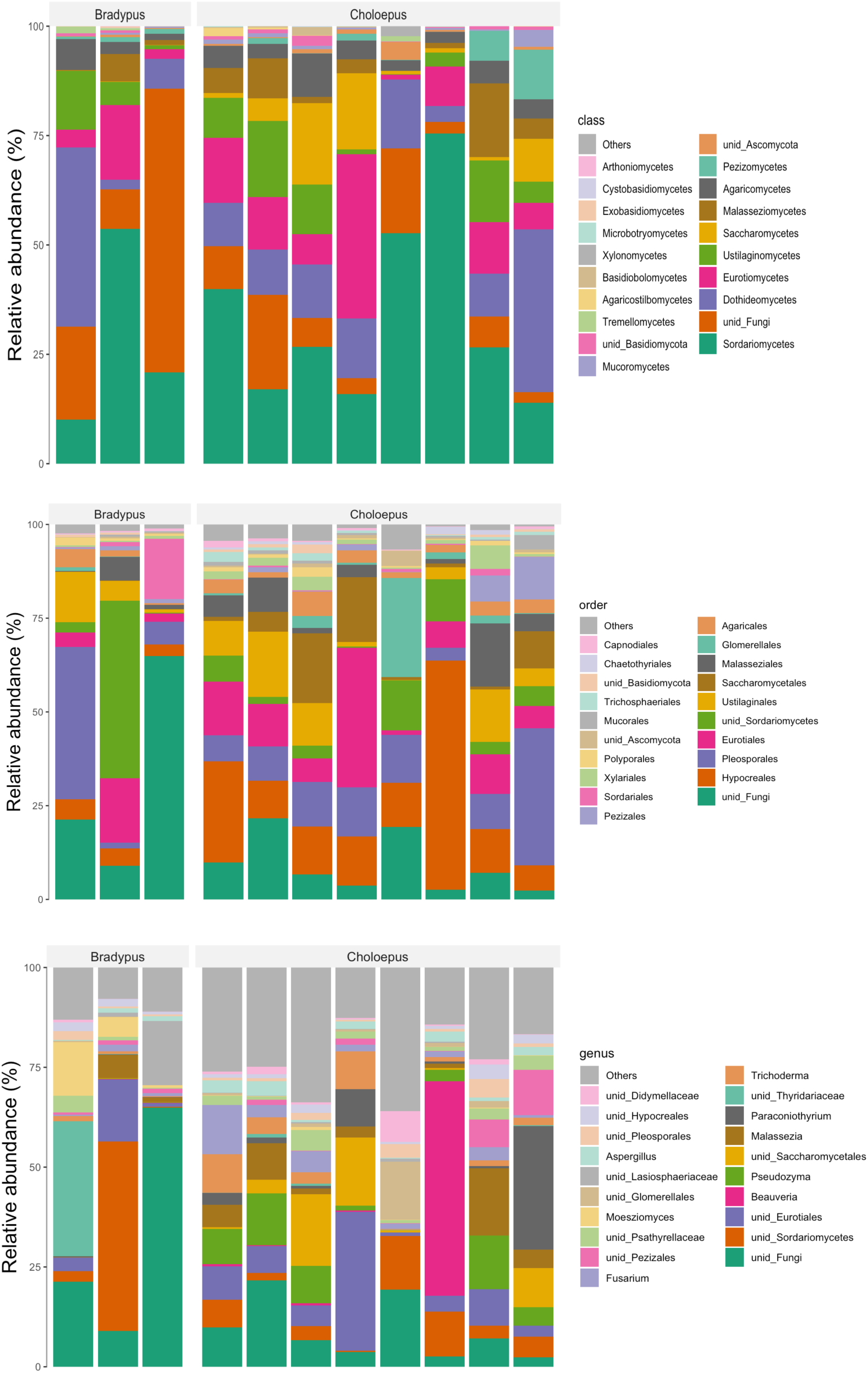
Relative abundance (%) of fungal class, order, and genus for ITS1 metabarcoding.

**Figure S6.**
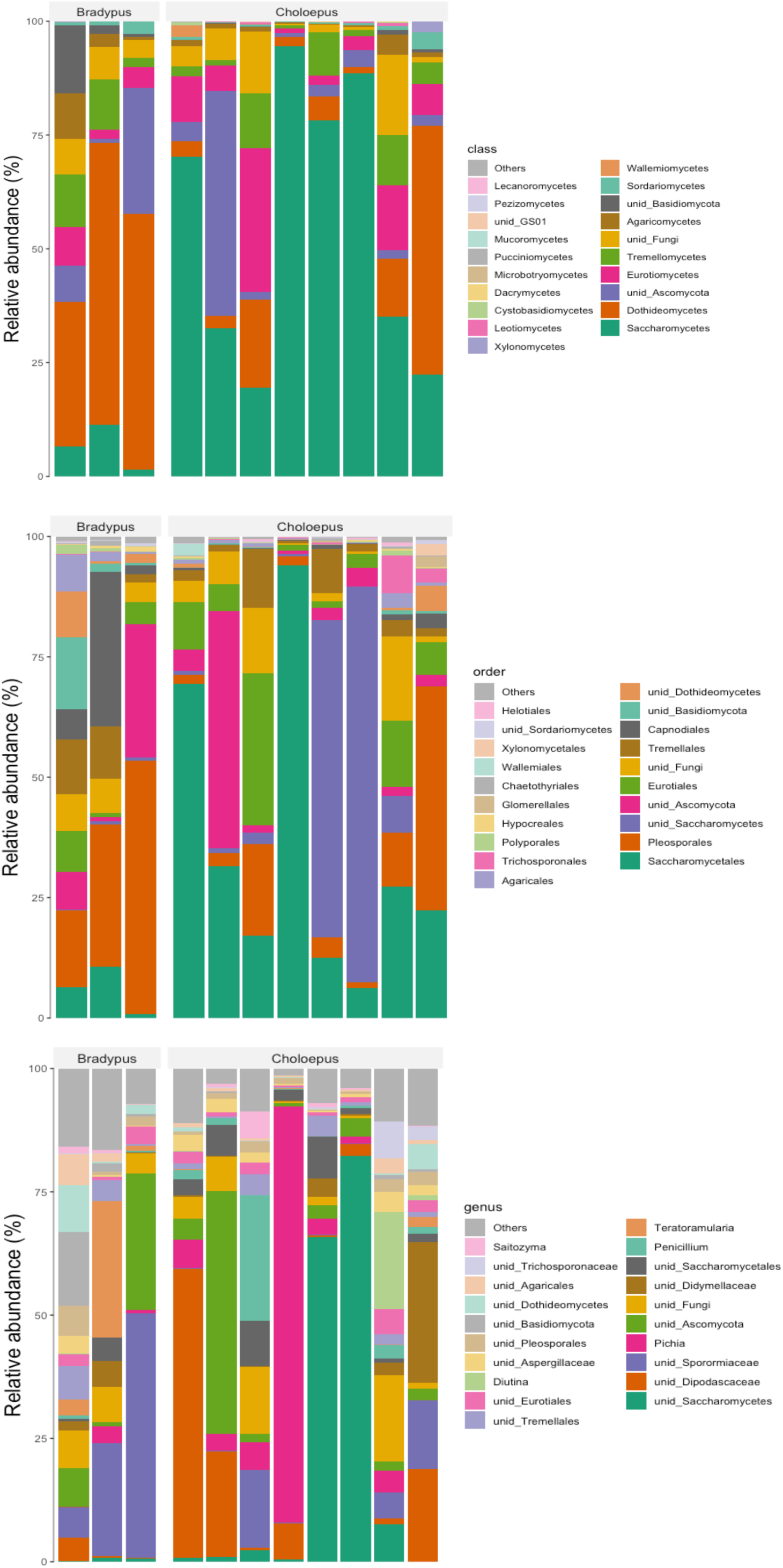
Relative abundance (%) of fungal class, order, and genus for ITS2 metabarcoding

**Figure S7.**
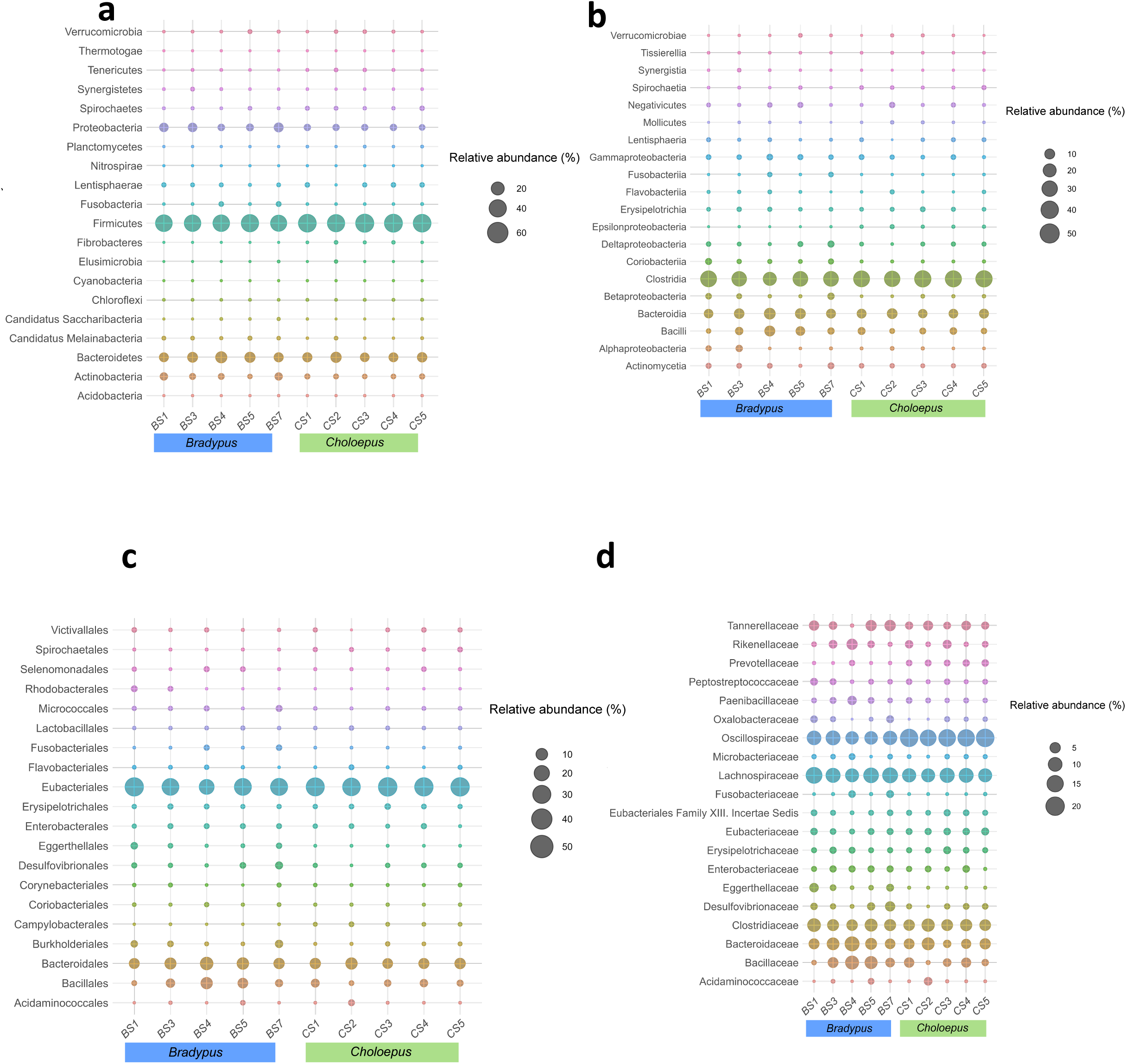
Relative abundances (%) of bacterial taxa per sample and species of sloth. Relative abundances presented here are based on the results from the search with Kaiju taxonomy classifier and metagenomic data. <u>Bacterial</u> phyla (**a**), <u>classes</u> (**b**), <u>orders</u> (**c**), and families (**d**) per sample are shown. Only the top 20 most abundant taxa are included. BS and CS represent samples from *B. variegatus* and *C. hoffmanni*, respectively. Unclassified reads are not included.

**Figure S8.**
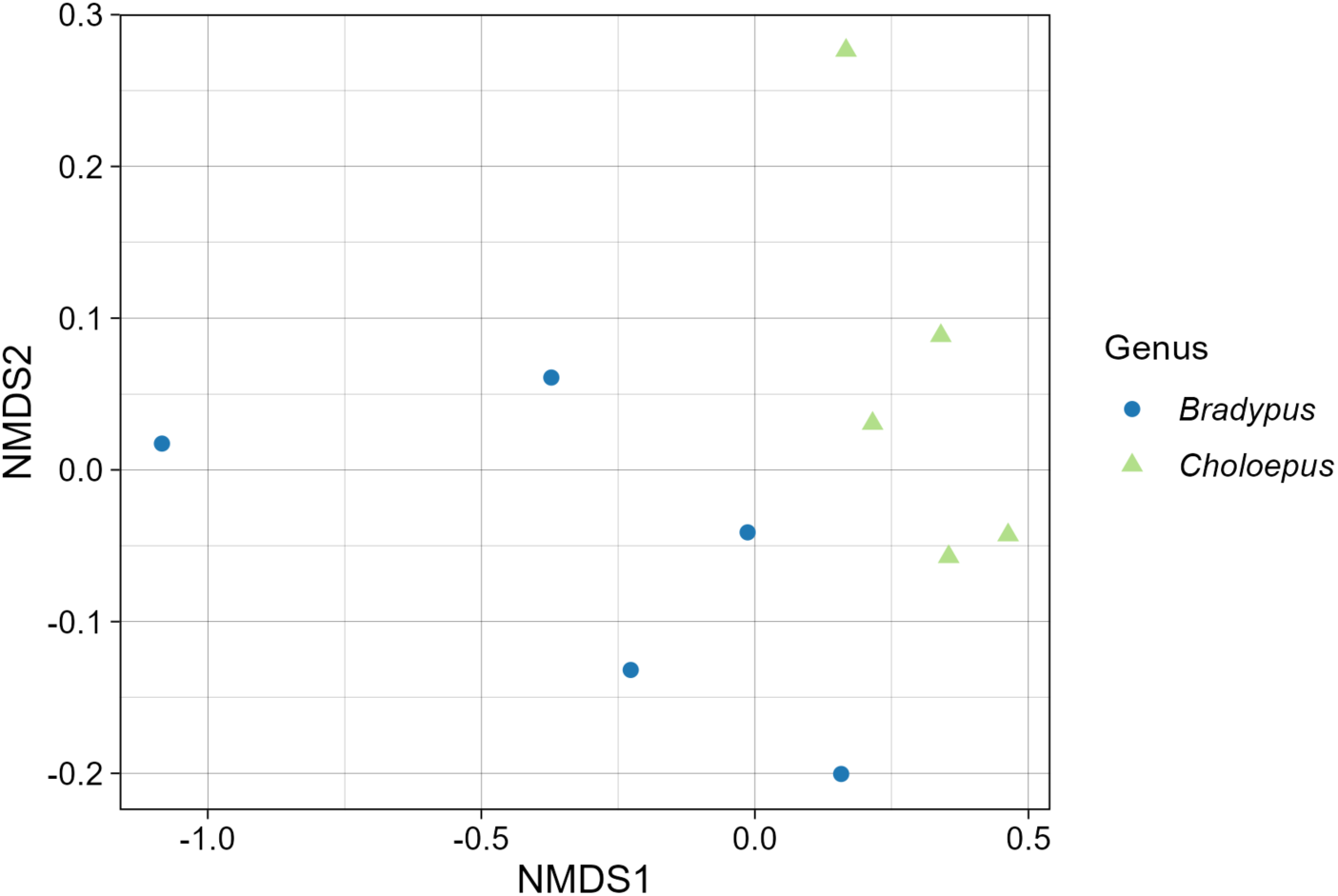
Nonmetric multidimensional scaling (NMDS) for the *Choloepus variegatus* and *Bradypus hoffmanni* overall microbial communities calculated with a Bray-Curtis distance matrix. Only shotgun metagenomic data was used. The final stress value for the 2D solution was 0.01508.

**Figure S9.**
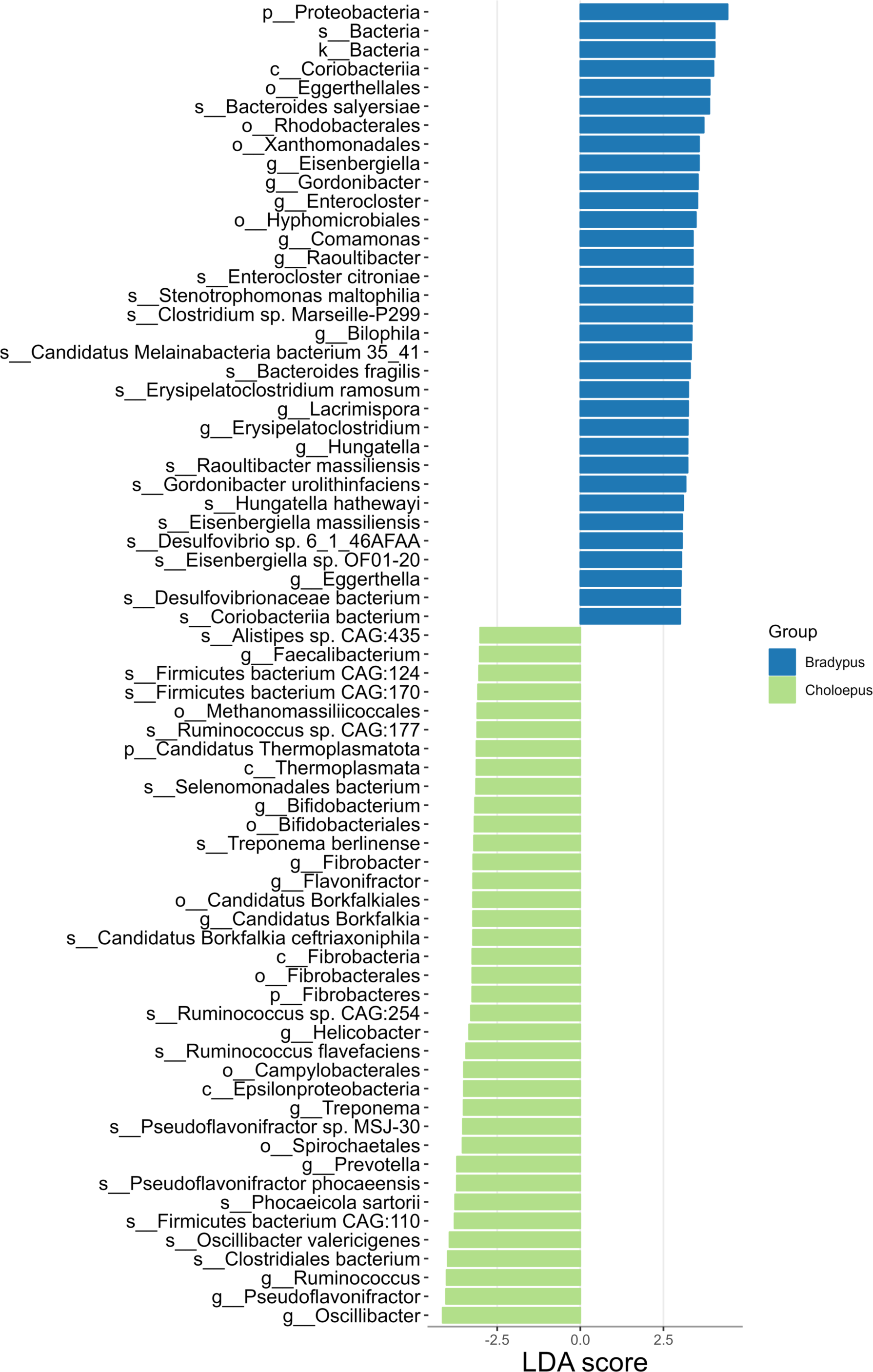
Linear Discriminant Analysis (LDA) scores in LEfSe for the *Choloepus variegatus* and *Bradypus hoffmanni* microbial communities based on the presence and absence from metagenomic data. The letters before the taxonomic ranks represent kingdom (k); phylum (p); class (c); order (o); genus (g); and species (s).

**Figure S10.**
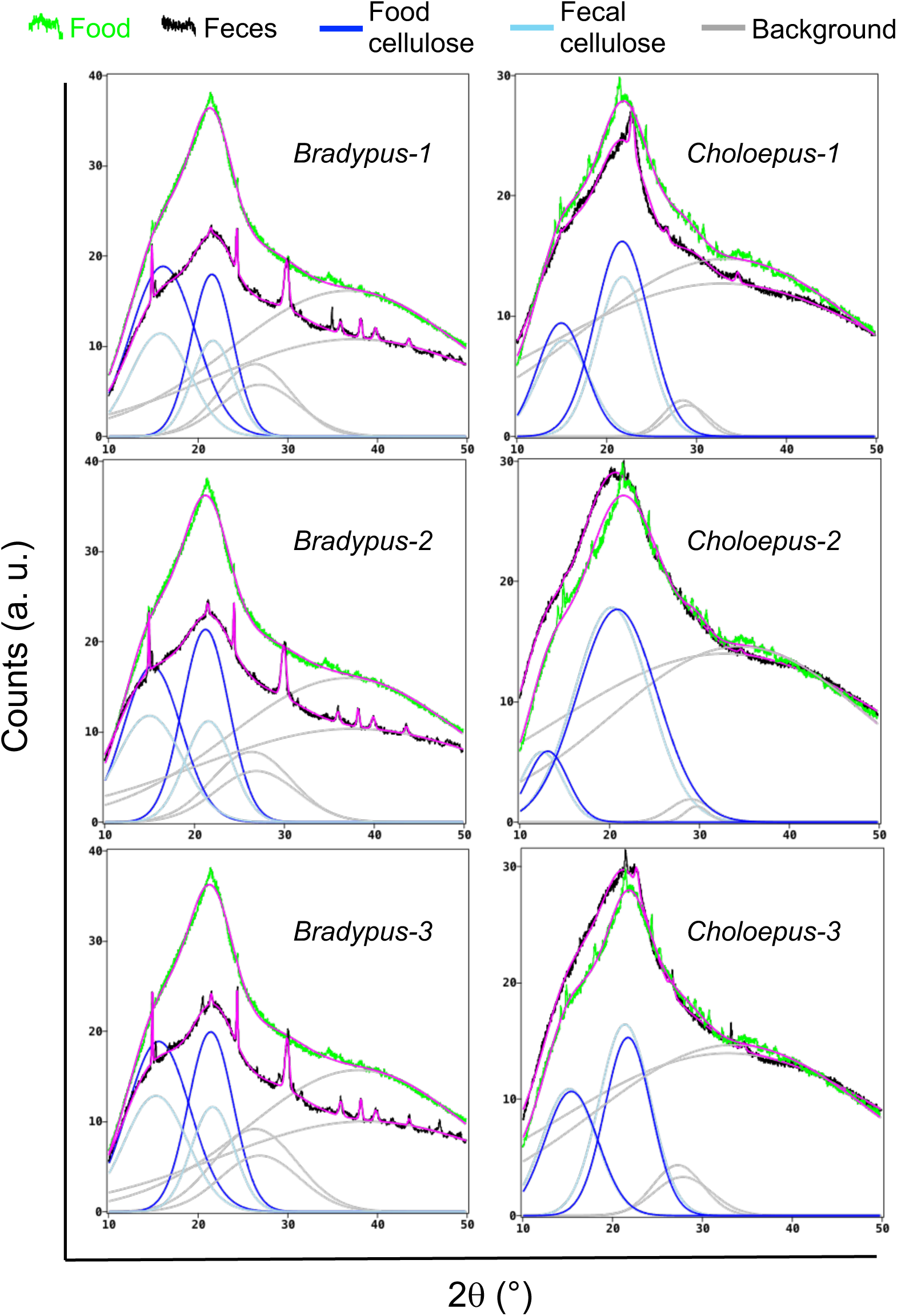
X-ray diffractograms for all three replicates of powders of food and the corresponding feces for sloth species *B. variegatus* (a) and *C. hoffmanni* (b). Green noisy curves correspond to the experimental signals of foods, black noisy curves correspond to experimental fecal signals. Blue and light blue curves are deconvoluted Gaussian functions assigned to cellulose in the food and feces, respectively. Gray curves represent background Gaussian functions. Magenta curves represent the fitted functions which are the sum of the background functions and the deconvoluted peaks assigned to cellulose. The sharp peaks present in the diffractogram of *B. variegatus* feces are due to calcium oxalate crystals (RRUFF Database, https://rruff.info/Whewellite/R050526); several Gaussian peaks were added to conveniently fit these signals but were excluded from the figure for the sake of simplicity. Choloepus-2 replicate was eliminated due to a negative value.

## Supplementary Tables

All supplementary tables and data are in https://github.com/pchaverri/sloths/

**Table S1**. These data come from running the Kaiju nr_euk database with assembled metagenomes.

**Table S2**. Pivot and interactive table with taxonomy and abundance, from metagenomic sequencing, both sloth species, and Kaiju nr_euk database.

**Table S3**. These data come from running the Kaiju fungi database with assembled metagenomes.

**Table S4**. Results of Kraken database search for *Bradypus*.

**Table S5**. Results of Kraken database search for *Choloepus*.

**Table S6**. These data come from running Kaiju nr_euk database with an assembled metatranscriptome of a *Choloepus hoffmanni* RNA pool.

**Table S7**. Pivot and interactive table with taxonomy and abundance, from metatranscriptomic sequencing, *Choloepus* pool, and Kaiju nr_euk database (from Table S6).

**Table S8**. Abundance (#reads) and taxonomy of ITS1 fungi metabarcoding.

**Table S9**. Abundance (#reads) and taxonomy of ITS2 fungi metabarcoding.

**Table S10**. Raw eggNOG-mapper results for *Bradypus variegatus*, all samples combined, from metagenomic data.

**Table S11**. Raw eggNOG-mapper results for *Choloepus hoffmanni*, all samples combined, from metagenomic data. Due to the large size of this file, the table has been divided into 9 parts. Parts 1-8 include all classified COGs. Part 9 consists of unclassified COGs.

**Table S12**. Summary of eggNOG-mapper functional annotation for metagenomic data for *Choloepus* and *Bradypus*. Only fungi and bacteria are included (from Tables S10 and S11).

**Table S13**. Raw results from eggNOG mapper classification from *Choloepus* RNA pools.

**Table S14**. Pivot and interactive tables with data from eggNOG mapper classification from *Choloepus* RNA pools (Table S13).

In addition, **interactive Krona figures in HTML format for** Figure 3 are included (<u>Krona_Bradypus.html</u>) and (<u>Krona_Choloepus.html</u>). These HTML files include all taxa (Prokaryota, Eukaryota and Viruses).

### EXTENDED METHODS

#### Permits

All permits for sampling sloth feces were obtained from the Institutional Commission of Biodiversity of the University of Costa Rica (resolution N° 339) and Judy Avey-Arroyo, the owner and legal representative of the Sloth Sanctuary (https://www.slothsanctuary.com/).

#### Sloths’ diet

The diet of *Bradypus* is exclusive of *Cecropia* spp. leaves. Daily, the animals are fed with 400 grams of fresh *C. insignus* and*/*or *C. obtusifolia* leaves. On the other hand, the diet provided to *Choloepus* sloths is more complex given their herbivorous-omnivorous nature ^1^. These animals are fed daily with 110 g of carrots (*Daucus carota* subsp. *sativus*), 110 g of sweet potato (*Ipomoea batatas*), 110 g of green beans (*Phaseolus vulgaris*), 110 g of chayote squash (*Sechium edule*), 200 g of Chinese watercress (*Ipomoea aquatica*), 4 units of asparagus beans (*Vigna unguiculata* subsp. *sesquipedalis*), and 230 g of beach almond leaves (*Terminalia catappa*). Additionally, once a week they are provided with 110 g of green mango (*Mangifera indica*) or water apple (*Syzygium aqueum*). This diet was formulated by those in charge of the sloth sanctuary, considering their experience and observations for more than 30 years. Using this diet, no high glucose levels, kidney problems or other problems associated with the diet have been observed in sloths of the genus *Choloepus*.

#### Sample collection

Due to the small number of animals and low defecation frequency, fewer samples were obtained from *B. variegatus* than for *C. hoffmanni* (see next section). The samples for this study were taken in three sampling campaigns: March 17-21, 2022; January 30-February 2, 2023 and January 29-30, 2024. All fecal samples were collected within 1 hour after defecation. The samples for DNA extraction were collected in 50 ml Falcon tubes, which were transported on ice to the laboratory, and frozen at -80°C until processing. Samples for RNA extraction were collected in 50 ml Falcon tubes and immediately suspended in RNAlater® preservative (Sigma Aldrich, USA). The samples were transported on ice to the laboratory, and frozen at -80°C until processing.

#### DNA and RNA extractions, and sequencing

For ITS metabarcoding analysis, fecal samples from 3 *B. variegatus* and 8 *C. hoffmanni* individuals were used. For shotgun metagenomics fecal samples from 5 *B. variegatus* and 5 *C. hoffmanni* individuals were used. DNA was extracted using DNeasy PowerSoil kit (QIAGEN, Hilden, Germany), with modifications. Briefly, the starting fecal matter (250 mg) was added to a microcentrifuge tube (2 mL) containing a spatula tip of three types of powdered glass with different particle size (850, 500 and 250 μm) along with a ceramic bead. Then, C1 solution was added (1000 μL) and sample was homogenized using a FastPrep instrument (5 m/s, 30 s; MP Biomedicals, California, US). The samples were then centrifuged (15.000 × *g*, 8 min) and 650 µL of supernatant were transferred to a clean new tube (1.5 mL). C2 solution (250 µL) was added to the supernatant, vigorously mixed (5 s) and the tubes were incubated on ice (10 min), followed by centrifugation (15.000 × *g*, 1 min). The supernatant was transferred to a new tube (1.5 mL) and solution C3 was added (200 µL). After incubating on ice (10 min), the samples were centrifuged (10.000 × *g*, 10 min) and the supernatant was transferred to a clean tube (2 mL). Solution C4 was added to the supernatant and strongly mixed (5 s). A volume of 600 µL was loaded onto a EZ-10 spin column (Bio Basic®, Toronto, Canada) for each sample and centrifuged (15.000 × *g*, 1 min). The flow-through was discarded, and the previous step was repeated until the remaining supernatant was loaded on the column. Next, the column was placed into a clean collection tube (2 mL), washed twice with solution C5 (500 µL), and centrifuged (15.000 × *g*, 1 min) after each wash. Following the wash steps, the column was centrifuged (15.000 × *g*, 1 min) until dry. Ethanol 80% (500 µL) was added to the column, centrifuged (15.000 × *g*, 1 min) and the column was placed into a microcentrifuge tube (1.5 mL). Finally, solution C6 (70 µL) were added to the column, centrifuged (15.000 × *g*, 1 min) and the eluted DNA was stored at –80 °C for further use. To obtain a comprehensive microbiota profiling (e.g., prokaryotes, eukaryotes, and viruses), genomic DNA samples were sent to Novogene Inc. (Sacramento, CA, USA) for shotgun metagenome sequencing (Illumina NovaSeq PE150). To better characterize the fungal communities in the fecal samples, targeted amplicon metagenomics (metabarcoding) was also outsourced to Novogene (Illumina NovaSeq PE250). The Internal Transcribed Spacers of the nuclear ribosomal DNA (ITS nrDNA) region is considered the official barcode for fungi and thus, has the most comprehensive and curated databases ^2,3^. For metabarcoding, ITS1 and ITS2 were sequenced to recover as much fungal diversity as possible ^4^. The primers used were ITS1: IT5-1737F and ITS2-2043R; and ITS2: ITS3-2024F and ITS4-2409R. As part of the sequencing services, Novogene performed raw data filtering, which consisted of removing reads containing adapters and primers, N > 10%, and low-quality (Qscore ≤ 20) bases.

For metatranscriptomics analysis, a pool of 23 fecal samples from *C. hoffmanni* individuals were used. Unfortunately, not enough *Bradypus* fecal material was obtained to obtain RNA with the quantity and quality for metatranscriptome sequencing. RNA extraction and metatranscriptome sequencing were performed at the sequencing facility of the German Centre for Infection Research (DZIF), Associated Partner Site Charité (Berlin, Germany) using an Illumina NextSeq 550 system (150 cycles paired-end). Fecal samples from *Choloepus hoffmanii* kept in RNAlater buffer were transported to DZIF where total nucleic acids (RNA) were extracted using the MagNA Pure 96 DNA and Viral NA Large Volume Kit (Roche Diagnostics, Indianapolis, IN, USA) in a Magna Pure 96 Instrument (Roche Diagnostics), following the manufacturer’s protocols. The RNA extractions were pooled to have enough concentration and quality for sequencing. RNA libraries were prepared for the sample according to KAPA HyperPrep manufacturer protocol (Roche Diagnostics).

#### Bioinformatic analyses

##### Metagenomics, metatranscriptomics, and functional annotation

All bioinformatic analyses were run on the Kabré supercomputer (CNCA-CONARE, Costa Rica). Once the raw .fastq files were obtained from Novogene and DZIF, Seqtk v.1.4 (https://github.com/lh3/seqtk/) and BBDuk v.38.84 (https://sourceforge.net/projects/bbmap/), respectively, were used to trim and filter low-quality reads, using default options (trim adapters, trim low quality reads <Q20, and discard short reads <20 bp). To remove host reads from the trimmed files, two available sloth genomes (*Bradypus variegatus* GCA_004027775.1 and *Choloepus hoffmanni* GCA_000164785.2) were first downloaded from GenBank and then created separate databases with those genomes using Bowtie2 v.2.2.9 ^5^. Following, the metagenome reads (forward R1 and reverse R2) were mapped to the host databases using Bowtie2 (“very-sensitive” mode). The resulting .sam files were converted to .bam format using SAMtools v.1.9 ^6^. The unmapped reads (i.e., without the host reads) were first extracted and then sorted according to read name with SAMtools. The reads were then divided into R1 and R2 and converted to .fastq files with BEDtools v.2.27.1 ^7^. The assembly of the metagenomes or metatranscriptomes was done with SPAdes v.3.14.1 (“meta” or “rna”) ^8–10^. EggNOG-mapper v.2.1.6 ^11^ was used for functional annotation of the assembled metagenome and metatranscriptome reads. EggNOG-mapper predicts orthologous groups and phylogenies from the eggNOG database (http://eggnog5.embl.de) to then transfer functional information from those predicted orthologous groups ^11^. Contigs and transcripts less than 500 bp were removed. EggNOG-mapper was run using MMseqs as the search step algorithm ^12^. The .fastq files with host reads removed have been deposited in GenBank under BioProject PRJNA991536.

Two tools to taxonomically classify the reads, i.e., Kaiju v.1.9.2 ^13^ and Kraken2 ^14^, were utilized and compared. For this, only the assembled contigs/scaffolds and transcripts that were longer than 500 bp were used. Kaiju has been reported as equally sensitive to Kraken2, but Kaiju is the only classifier that includes fungal and other eukaryotes databases from the NCBI RefSeq database and a protein-based classifier ^13,15,16^. For Kaiju, the analysis was first run using “nr_euk” database v.2022-03-10, which includes prokaryotes, viruses, and eukaryotes. Because with nr_euk significantly fewer reads were obtained that matched fungi, the analysis was also run using only the fungi database v.2023-05-06. The analyses were run for each sample separately. For Kraken2, the standard database (v.2023-4-25), plus fungi and protozoans, was used. The results shown in the present study are based on Kaiju nr_euk database because when the search was run with the Kaiju fungi and Kraken2 databases, many fungal groups were missing (e.g., Neocallimastigomycota from Kaiju fungi, and most non-Dikarya from Kraken2) (supplementary Table S1). Relative abundance (%) calculations and bar graphs were done in the package microeco v.1.1.0 ^17^ in RStudio v.2023.03.1+446. All the resulting and supplementary data are deposited in the public repository GitHub https://github.com/pchaverri/sloths.

##### Targeted amplicon metagenomics/metabarcoding of fungi

DADA2 v.1.26.0 ^18^ implemented in RStudio was utilized for processing raw sequencing data. This included quality inspection, filtering, trimming, the merging of paired-end reads, inference of Amplicon Sequence Variants (ASVs) ^19^, and subsequent removal of chimeras (adapters and primers were already removed by Novogene Inc.). Quality control involved filtering and trimming forward and reverse sequences with a quality score threshold of < 20, alongside a maximum expected errors (maxEE) parameter set to 2. Subsequently, ASVs were clustered and subjected to chimera removal. Taxonomic assignments were done by performing similarity searches against the curated and quality-checked UNITE database v.2021 ^3,20^ using the DECIPHER package v.2.0 ^21^. Relative abundance (%) calculations and bar graphs were also done in microeco. The raw .fastq files have been deposited in GenBank under BioProject PRJNA991536. Additional results and supplementary data are deposited in GitHub public repository https://github.com/pchaverri/sloths.

##### Community analysis

The microbial communities of *B. variegatus* and *C. hoffmanni* were compared using shotgun metagenomic sequencing results. Two datasets were analyzed: the first dataset included all Kingdoms matching the ASVs, while the second consisted only of reads that matched fungi to detect Neocallimastigomycota. For beta diversity analysis, the Hellinger transformation was applied to the ASV counts table. The "adonis" and "permutest" functions from the Vegan package^22^ were used to evaluate differences in the microbial communities between sloth species and homogeneity across samples, with permutation parameters set to 5,000 for both tests. Visualization was performed using Non-metric Multidimensional Scaling (NMDS) in Phyloseq from Bioconductor ^23^. For this, count data was transformed into a Bray-Curtis distance matrix, and the analysis was conducted with a set seed. Linear Discriminant Analysis (LDA) diagrams were created using the microeco package. Relative abundance, NMDS, LDA plots, and all analyses were performed using R v.4.3.3.

##### Comparative analysis of cellulose degradation using X-ray diffraction (XRD)

XRD was selected as the method for determining plant biomass degradation due to its ability to provide detailed structural information, quantify crystallinity, and monitor changes in a non-destructive and sensitive manner^24^. XRD of powder samples was used to quantitatively assess crystalline and amorphous components present in materials ^25,26^. The areas or intensities of deconvoluted signals in diffractograms assigned to a specific substance are considered proportional to the content of the substance. In the case of samples containing cellulose, the relative contents of crystalline and amorphous cellulose have been previously assessed with this deconvolution technique ^27,28^. Three dried and ground samples of each sloth’s specific feed were analyzed by XRD at 2θ values between 10° and 50°, where signals associated to cellulose appear^29^. Identical procedures of sample preparation and XRD measurements were applied to all samples.

Sloth food samples (*C. peltata* leaves for *Bradypus* and mixture for *Choloepus*) and feces from both sloth species were dried at 50 °C for 120 h. Once dried, each sample of food and feces were ground with a blender to obtain fine powders. The sample of food mixture for *Choloepus* was prepared with amounts of each ingredient proportional to specified quantities indicated by the sloth keepers (see above). Samples were analyzed with a Bruker D8 Advance ECO X-ray powder diffraction equipment with Bragg-Brentano geometry; under the following measurement conditions: power 1000 W, rotation 15 rpm, angles 2θ in the interval 10° - 50°, step size 0.02°, step time 8.0 s, and aperture of lens 1.5°. Each resulting diffractogram was deconvoluted by summing four Gaussian functions centered at 2θ values about 15°, 22°, 28° and 38°. The first two Gaussian peaks were assigned to cellulose ^29^, and the remaining were used to fit the background (signals assigned to other substances than cellulose, such as lignin, are considered as part of the background). The experimental diffractograms were fitted by non-linear regression using the software for symbolic and numeric computation Maple (Maplesoft, Waterloo, Canada). The areas of the peaks centered about 15° (𝐴15°) and 22° (𝐴22°) for the fecal samples were compared with the peak areas calculated for the corresponding food to estimate the cellulose degradation ratio for both species of sloths:

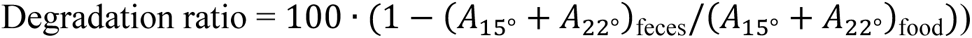

